# MLL3 regulates the *CDKN2A* tumor suppressor locus in liver cancer

**DOI:** 10.1101/2022.06.07.495174

**Authors:** Yadira M. Soto-Feliciano, Changyu Zhu, John P. Morris, Chun-Hao Huang, Richard P. Koche, Yu-jui Ho, Ana Banito, Chun-Wei Chen, Aditya Shroff, Sha Tian, Geulah Livshits, Chi-Chao Chen, Myles Fennell, Scott A. Armstrong, C. David Allis, Darjus F. Tschaharganeh, Scott W. Lowe

## Abstract

Mutations in genes encoding components of chromatin modifying and remodeling complexes are among the most frequently observed somatic events in human cancers. For example, missense and nonsense mutations targeting the mixed lineage leukemia family member 3 (*MLL3*/*KMT2C*) histone methyltransferase occur in a range of solid tumors and heterozygous deletions encompassing *MLL3* occur in a subset of aggressive leukemias. Although *MLL3* loss can promote tumorigenesis in mice, the molecular targets and biological processes by which MLL3 suppresses tumorigenesis remain poorly characterized. Here we combined genetic, epigenomic, and animal modeling approaches to demonstrate that one of the mechanisms by which MLL3 links chromatin remodeling to tumor suppression is by co-activating the *Cdkn2a* tumor suppressor locus. Disruption of *Mll3* cooperates with *Myc* overexpression in the development of murine hepatocellular carcinoma (HCC), in which MLL3 binding to the *Cdkn2a* locus is blunted, resulting in reduced H3K4 methylation and low expression levels of the locus-encoded genes, *Ink4a* and *Arf*. Conversely, elevated *MLL3* expression increases its binding to the *CDKN2A* locus and co-activates gene transcription. Endogenous *Mll3* restoration reverses these chromatin and transcriptional effects and triggers *Ink4a*/*Arf*-dependent apoptosis. Underscoring the human relevance of this epistasis, we found that genomic alterations in *MLL3* and *CDKN2A* display mutual exclusivity in human HCC samples. These results collectively point to a new mechanism for disrupting *CDKN2A* activity during cancer development and, in doing so, link MLL3 to an established tumor suppressor network.

## INTRODUCTION

Hepatocellular carcinoma (HCC) is a deadly primary liver cancer with a 5-year survival rate of only 18% (Jemal et al., 2017). HCC is currently the fourth most frequent cause of cancer-related mortality worldwide with a continuously growing incidence (Llovet et al., 2021). Genomic alterations found in HCC are highly diverse, and are characterized by *TERT* (telomerase reverse transcriptase) promoter mutations, amplifications or chromosomal gains encompassing the *MYC* oncogene, activating hotspot mutations in *CTNNB1* (β-catenin), and inactivating mutations and deletions in the *TP53* and *CDKN2A* tumor suppressor genes (Schulze et al., 2015; The Cancer Genome Atlas Research Network, David A. Wheeler, Lewis R. Roberts, 2017). Beyond these well-studied drivers, HCC frequently harbors mutations in one or more chromatin modifying enzymes, including *MLL3* (*KMT2C*) (Fujimoto et al., 2012; Kan et al., 2013).

MLL3 is a component of the COMPASS-like complex that has structural and functional similarities to the developmentally essential *Drosophila* Trithorax-related complex (Schuettengruber et al., 2017). This multiprotein complex controls gene expression through its histone H3 lysine 4 (H3K4) methyltransferase activity that establishes chromatin modifications most often associated with transcriptional activation (Shilatifard, 2012). Interestingly, mutations in another homologous H3K4 methyltransferase of the complex, *MLL4* (*KMT2D*) are also found in HCC (Cleary et al., 2013). *KDM6A* (*UTX*), a H3K27 demethylase within the COMPASS-like complex, has been recently identified as a potent tumor suppressor in liver and pancreatic cancers (Revia et al., 2021). While these observations suggest that epigenetic-based mechanisms of gene regulation controlled by the MLL3 complex can constrain HCC development, the molecular targets and biological processes that underlie MLL3’s tumor suppressor activities remain poorly understood.

## RESULTS

### *Mll3* is a tumor suppressor in *Myc*-driven liver cancer

To determine whether *Mll3* loss impacts liver cancer development, we applied hydrodynamic tail vein injection (HTVI) in wild-type mice to directly introduce genetic manipulations into a subset of adult hepatocytes *in vivo* (Bell et al., 2007). This gene delivery method facilitates the study of oncogene-tumor suppressor interactions by combining stable genomic integration of oncogenic cDNAs (transposon vector) and transient expression of plasmids encoding Cas9 and single guide RNAs (sgRNAs) to disrupt tumor suppressor genes (Largaespada, 2009; Moon et al., 2019; Tschaharganeh et al., 2014; Xue et al., 2014). Since *MLL3* mutations co-occur with *MYC* genomic gains and amplifications in human HCC tumors (**Fig. 1A**), we used the HTVI approach to test whether disruption of *Mll3* by CRISPR could cooperate with *Myc* overexpression to drive murine liver cancer (**Fig. 1B**).

**Figure 1.**
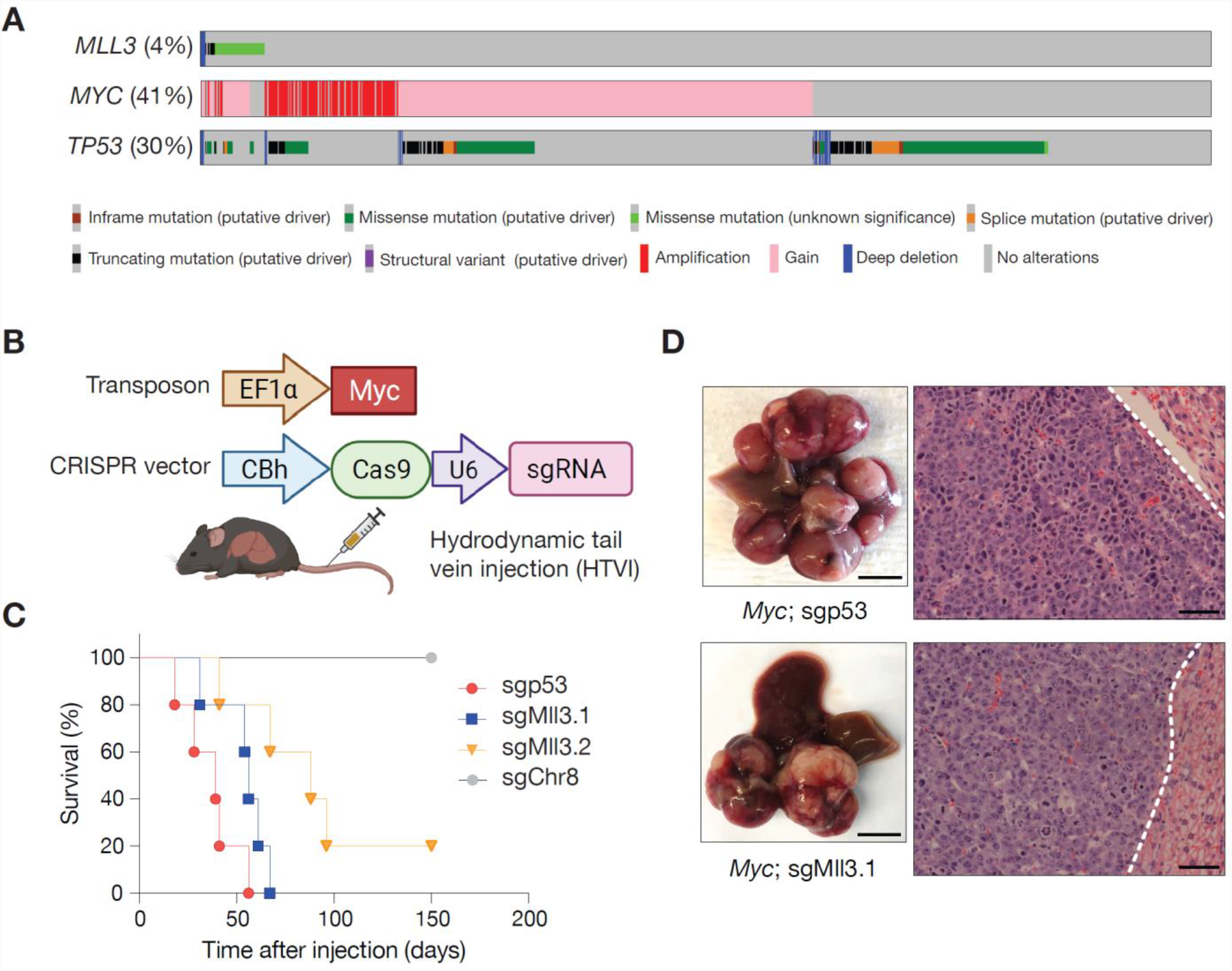
*Mll3* constrains *Myc*-driven tumorigenesis. **(A)** Oncoprints displaying genomic mutations and deletions of *MLL3* and *TP53*, as well as gains and amplifications of *MYC* in merged publicly available datasets (TCGA ((Schulze et al., 2015; The Cancer Genome Atlas Research Network, David A. Wheeler, Lewis R. Roberts, 2017)), MSK (Harding et al., 2019; Zheng et al., 2018), INSERM (Schulze et al., 2015), and RIKEN (Fujimoto et al., 2012)) of 1158 sequenced hepatocellular carcinomas (HCCs). **(B)** Schematic for hydrodynamic tail vein injection (HTVI) of gene delivery into murine livers. Vectors permitting stable expression of *Myc* transposon (top) and transient expression of Cas9 and sgRNAs targeting putative tumor suppressors (bottom) via sleeping beauty transposase were introduced into hepatocytes by HTVI. **(C)** Survival curve of mice injected with *Myc* transposon and pX330 expressing two independent sgRNAs targeting *Mll3* after HTVI injection (*Myc;* sgMll3.1, n = 5; *Myc;* sgMll3.2, n = 5). *Myc;* sgp53 (n = 5), and *Myc;* sgChr8 (n = 5) serve as controls. (**D)** Representative images (left, liver macro-dissection, scale bar: 0.5 cm; right, H&E staining, scale bar: 100 μm) of mouse liver tumors generated by HTVI delivery of *Myc* transposon and *in vivo* gene editing (top, *Myc*; sgp53; bottom, *Myc*; sgMll3.1). The dash lines indicate the boundaries between liver tumors and non-tumor liver tissues.

Mice injected with a *Myc* cDNA transposon and two independent Cas9/*Mll3* sgRNAs (*Myc*; sgMll3.1 or *Myc*; sgMll3.2) combinations developed liver tumors with a slightly later onset compared to mice receiving an sgRNA targeting the tumor suppressor *Trp53* (*Myc*; sgp53) (**Fig. 1C-D**). Adult livers injected with *Myc* and a control sgRNA (sgChr8) did not succumb to disease over the observation period (**Fig. 1C**). These findings were confirmed in a second independent cohort of mice (**Fig. S1A**). Analyses of tumor-derived genomic DNA revealed insertions and deletions (indels) in either *Mll3* or *Trp53* (hereafter simply referred to as *p53*) depending on the genotype of tumor-derived cells (**Fig. S1B**). DNA sequencing of the CRISPR-targeted region from two independent *Myc*; sgMll3 tumors revealed either heterozygous or homozygous indels predicted to generate premature stop codons (**Fig. S1C**). These data imply that even partial suppression of *Mll3* can promote tumorigenesis. In support of this, GFP-linked *Mll3* shRNAs efficiently cooperated with *Myc* overexpression to drive liver cancer producing tumors with 50-80% reduction in *Mll3* mRNA expression (**Fig. S1D-S1G**). shMll3.2 resulted in less potent knockdown than shMll3.1, yet produced faster tumor formation, suggesting that similar to acute myeloid leukemia (Chen et al., 2014), *Mll3* can likely act as a haploinsufficient tumor suppressor in liver cancer (**Fig. S1E and S1G**).

### *Mll3* loss alters the chromatin landscape of liver cancer cells

MLL3 and MLL4 are histone methyltransferases that can deposit the H3K4 mono-methylation mark at genomic enhancers and intergenic regions during organ development (Hu et al., 2013). However, more recent studies indicate that MLL3 and MLL4 are also capable of binding to promoter regions (Cheng et al., 2014; Dhar et al., 2016; Wang et al., 2010), especially in the context of cancer (Soto-Feliciano et al., 2021). To determine the genomic binding patterns of MLL3 in HCC, we performed MLL3 ChIP-sequencing (ChIP-Seq) analysis in *Myc*; sgMll3 (sgMll3.1 which generates heterozygous or homozygous indels) and *Myc*; sgp53 liver cancer cell lines. Compared to sgp53 cells, *Mll3* deficiency resulted in a marked reduction in MLL3 chromatin binding at a subset of genomic loci (**Fig. 2A**). Approximately 40% of the peaks that were selectively lost in *Mll3*-deficient cells occurred at promoter regions, whereas unchanged MLL3 peaks between the two genotypes were more likely to be within intergenic regions (**Fig. 2B, Fig. S2**). Of note, the residual signal observed in ChIP-Seq most likely reflects the binding of MLL4 and/or remnant MLL3, since the antibody used in these experiments can recognize both MLL3 and MLL4 proteins (Dorighi et al., 2017). Our data suggest that, beyond the canonical action of MLL3 at gene enhancers, MLL3 and MLL4 can also occupy promoter regions in *Myc*-induced liver cancer.

**Figure 2.**
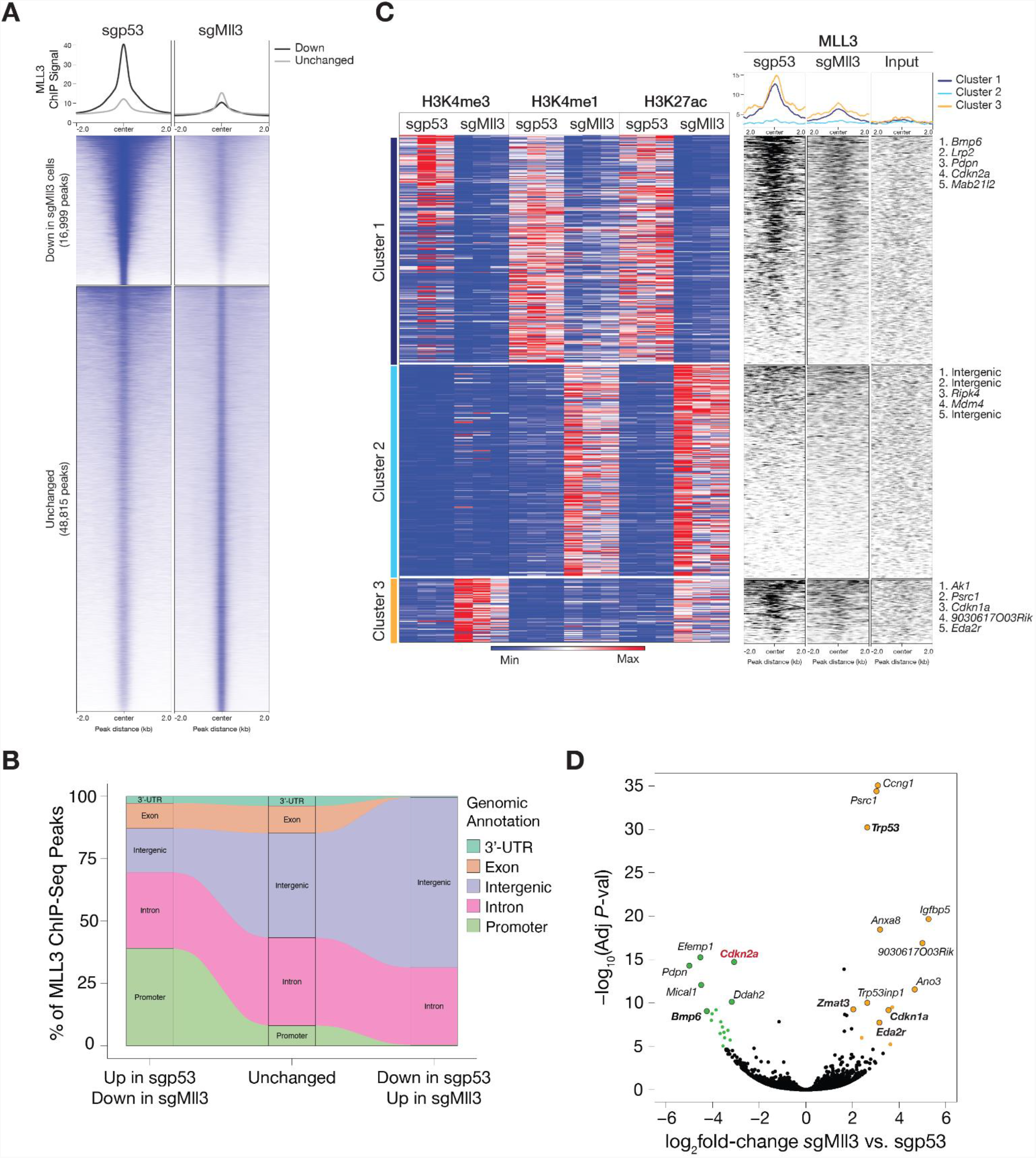
*Mll3* disruption alters the chromatin and transcriptional landscape of liver cancer cells. **(A)** Tornado plots showing MLL3 ChIP-Sequencing (ChIP-Seq) signal (peaks) that were down or remained unchanged in *Myc*; sgMll3 cells relative to *Myc*; sgp53 cells. **(B)** Alluvial plot showing the percentages of MLL3 ChIP-Seq peaks in different genomic elements in *Myc*; sgMll3 vs. *Myc*; sgp53 cells, including 16,999 peaks down, 48,815 peaks unchanged, and 265 peaks up in sgMll3 cells. Promoter regions were defined as transcriptional start sites (TSS) ± 2kb. **(C)** Heatmaps of histone modification ChIP-Seq signals (H3K4me3, H3K4me1, H3K27ac, left panel) and MLL3 ChIP-Seq signal (right panel) at promoter or intergenic regions in 3 independent *Myc;* sgMll3 and *Myc;* sgp53 liver tumor-derived cell lines. Cluster 1: loss of promoter and enhancer activity (loss of H3K4me3, H3K4me1, and H3K27ac); cluster 2: gain of enhancer activity (gain of H3K4me1 and H3K27ac); and cluster 3: gain of promoter activity (increase of H3K4me3). **(D)** Volcano plot of differentially expressed genes revealed by RNA-Sequencing of three independent *Myc; sg*Mll3 and *Myc; sg*p53 HCC cell lines. Genes in sgMll3 cells with more than 2-fold expression change and exceeding adjusted *P*-value<10^−5^ are color-labeled (orange: up-regulated; green: down-regulated). Some differentially expressed genes are labeled with gene symbols, and p53 targets are bolded.

Similar to the *Drosophila* Trithorax-related complex (Schuettengruber et al., 2017), the mammalian MLL3 and MLL4 complexes facilitate gene transcription by establishing permissive modifications on histone H3K4 via the MLL3 and MLL4 methyltransferases (Shilatifard, 2012). While MLL3 can establish H3K4 mono-methylation at enhancers in a redundant fashion with MLL4 (Shilatifard, 2012), *Mll3* inactivation also decreases H3K4me3 levels at the promoters of metabolism-related genes in normal murine livers (Valekunja et al., 2013) and human liver cancer cells (Ananthanarayanan et al., 2011). To determine whether *Mll3* haploinsufficiency impacts the local or global chromatin landscape of HCC cells, we performed ChIP-Seq analyses for H3K4 methylation and H3K27 acetylation in three independently derived *Myc*; sgMll3 and *Myc*; sgp53 tumor cell lines (**Fig. 2C**). Correlation analyses revealed three distinct clusters that include areas of enrichment and depletion for each tested histone modification between *Myc*; sgMll3 and *Myc*; sgp53 tumor cells (**Fig. 2C, Fig. S3A-C**). Loci associated with cluster 1 (reduced H3K4me3, H3K4me1, and H3K27ac in sgMll3 cells) showed the most pronounced and dynamic differences in chromatin modifications between the two liver tumor genotypes. In contrast, the loci in cluster 2 showed increased H3K4me1 and H3K27ac marks and most of them mapped to the intergenic regions of the genome, and cluster 3 loci with enhanced H3K4me3 likely represent some p53 target genes such as *Cdkn1a* and *Eda2r*.

To determine whether *Mll3* disruption is associated with these drastic changes in the chromatin landscape, we integrated our MLL3 ChIP-Seq from the *Myc*; sgp53 vs *Myc*; sgMll3 tumor cells with the chromatin modifications analyses (**Fig. 2C**). Interestingly, loci present in cluster 1 that displayed the most substantial changes in histone modifications involved genes that showed high MLL3 enrichment in *Myc*; sgp53 cells compared to the *Myc*; sgMll3 genotype. These data support a model whereby MLL3 binding to these loci facilitates the acquisition of a chromatin environment conducive for active gene transcription.

We next determined the output of these chromatin landscape changes by transcriptional profiling of the same set of *Myc*; sgp53 and *Myc*; sgMll3 liver cancer cell lines described above. Despite the broad binding of MLL3 across the genome, only 132 differentially expressed genes (DEGs) were significantly up-regulated (*P*<0.05, log_2_ fold-change>2) and 116 DEGs significantly down-regulated (*P*<0.05, log_2_ fold-change<-2) in *Myc*; sgMll3 liver tumor cells compared to *Myc*; sgp53 controls. As predicted, transcripts encoding p53 and p53 target genes such as *Ccng1, Cdkn1a*, and *Zmat3* (Bieging-Rolett et al., 2020) were elevated in *Myc*; sgMll3 cells, consistent with nonsense-mediated decay of truncated p53 transcripts and a concomitant reduction in p53 effector genes. Strikingly, some of the downregulated genes in *Myc*; sgMll3 lines mapped to loci enriched in cluster 1, including *Cdkn2a, Bmp6*, and *Lrp2* (**Fig. 2C-D, Fig. S3D-E**). In principle, cluster 1 genes that display: 1) MLL3 binding enrichment, 2) a histone profile associated with gene activation, and 3) reduced transcript levels in *Myc*; sgMll3 tumor cells should include genes that mediate the molecular and cellular effects downstream of *Mll3* disruption during tumor formation.

### Genomic inactivations of *MLL3* and *CDKN2A* display mutual exclusivity in human HCC

One genomic locus that stood out in our integrative analysis was *Cdkn2a*, which encodes for the *p16*^*Ink4a*^ and *p19*^*Arf*^ (*p14*^*ARF*^ in humans) tumor suppressor genes (hereafter referred to as *Ink4a* and *Arf*, respectively) (Gil and Peters, 2006). *CDKN2A* is located on the human chromosome 9p, and is deleted or epigenetically silenced in many cancer types (Sherr, 2012) including HCC (The Cancer Genome Atlas Research Network, David A. Wheeler, Lewis R. Roberts, 2017). While MLL3 likely regulates a plethora of genes that contribute to its tumor suppressive potential, the well-defined and potent anti-tumor functions of *Cdkn2a*-encoded proteins make them attractive candidates as functionally relevant MLL3 effectors.

To gain further insights into the potential relationship between *MLL3* and *CDKN2A*, we first looked at human cancer datasets to assess whether genomic alterations of *MLL3* and *CDKN2A* display an epistatic relationship that would be indicative of functional redundancy. Consistent with this possibility, an analysis of 1158 HCC samples derived from five independent, publicly available HCC sequencing datasets revealed a mutually exclusive relationship between the presence of *MLL3* and *CDKN2A* inactivating alterations including truncating mutations and deep deletions (**Fig. 3A**). Of note, our analyses excluded missense mutations owing to their unknown impact on MLL3 functions. Further dissection of transcriptional profiling datasets from human and mouse HCCs harboring known gene alterations using gene set enrichment analysis (GSEA) revealed that human tumors with *CDKN2A* deletions transcriptionally resemble both mouse and human HCC harboring *MLL3* alterations (**Fig. 3B-C**), but not those harboring *RB1* loss (**Fig. S4A**). Consistent with a degree of context-dependence that is observed for other chromatin regulators such as EZH2 (Ntziachristos et al., 2012; Soto-Feliciano et al., 2017; Souroullas et al., 2016; Tirode et al., 2014), *MLL3* and *CDKN2A* alterations displayed co-occurrence in several tumor types (**Fig. S4B**). Nevertheless, an overall analysis of a pan-cancer cohort showed a pattern of mutual exclusivity (Zehir et al., 2017) (**Fig. S4C**), with trends noted in ten individual blood and epithelial cancer types (**Fig. S4D**). While we cannot rule out the possibility that other factors drive these associations, our results support a biologically meaningful relationship between MLL3 and CDKN2A.

**Figure 3.**
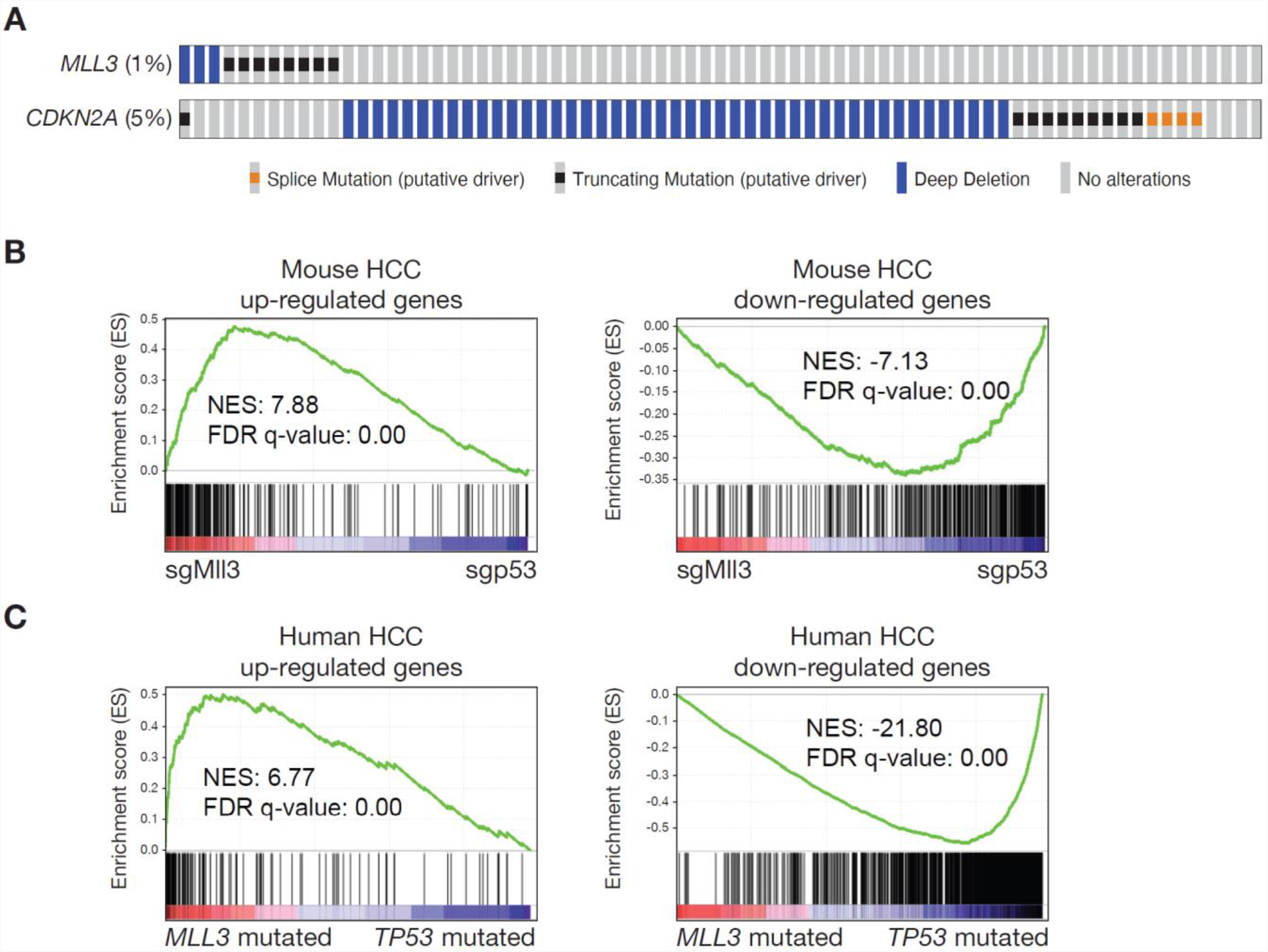
Genetic alterations in *CDKN2A* and *MLL3* show mutually exclusive patterns in human HCC and other cancers. **(A)** Oncoprints displaying genomic truncating mutations and deletions of *MLL3* and *CDKN2A* in merged datasets (TCGA, MSK, INSERM, and RIKEN) of 1158 sequenced HCCs. **(B** and **C)** Gene set enrichment analysis (GSEA) plots of human HCC transcriptional signatures with *CDKN2A* mutations or homozygous deletions in **(B)** mouse HCCs (*Myc*; sgMll3 or *Myc*; sgp53) and **(C)** human HCCs with mutations and deletions of either *MLL3* or *TP53*. Normalized enrichment scores (NES) and False discovery rate (FDR) q-values were calculated by GSEA.

### *Cdkn2a* locus is a genomic and transcriptional target of MLL3 in liver cancer

To explore the relationship between MLL3 and *Cdkn2a* in more detail, we tested whether genes encoded by *Cdkn2a* were direct targets of MLL3-regulated transcription. Indeed, *Cdkn2a* is a cluster 1 locus that display: 1) significant reduction in expression in *Myc*; sgMll3 cancer cells, with both decreased 2) levels of H3K4me1/3 and H3K27ac and 3) MLL3 binding at the promoters of *Ink4a* and *Arf* in sgMll3 tumors compared to sgp53 cells (**Fig. 2C-D, Fig. 4A**). The differential expression of *Ink4a* and *Arf* were also confirmed by qPCR, immunoblotting, and ChIP-qPCR analyses on multiple *Myc*; sgMll3 and *Myc*; sgp53 liver cancer lines (**Fig. 4B, Fig. S5A-B**). These results imply that *Cdkn2a* locus is a genomic and transcriptional target of MLL3 in liver cancer cells.

**Figure 4.**
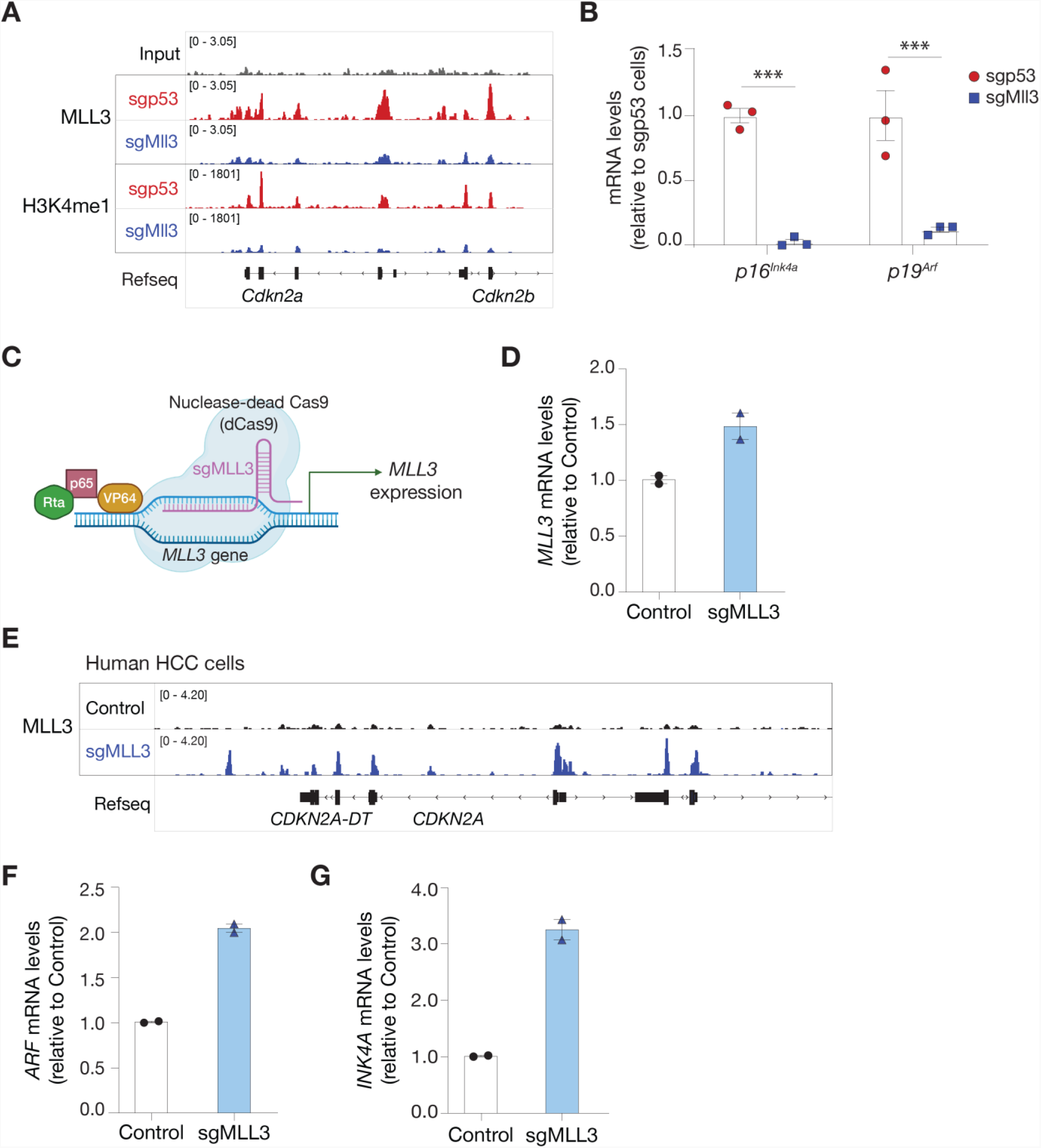
*CDKN2A* locus is a genomic and transcriptional target of MLL3 in liver cancer. **(A)** Genome browser tracks for MLL3 and H3K4me1 ChIP-Seq in *Myc;* sgp53 (red) and *Myc;* sgMll3 (blue) HCC cell lines at the *Cdkn2a* locus. **(B)** qPCR analysis for mRNA expression of *Ink4a* and *Arf* from 3 independent *Myc; sg*Mll3 and *Myc; sg*p53 HCC lines. Values are shown as mean ± SD. *** = *P*<0.001 (unpaired two-tailed t-test). **(C)** Schematic for CRISPR activation (CRISPRa) system of nuclease-dead Cas9 (dCas9) and VP64-p65-Rta (VPR) guided by sgMLL3 to activate *MLL3* expression in human HLE HCC cell line. **(D)** qPCR analysis for mRNA expression of *MLL3* in HLE cells with sgGFP (control) or sgMLL3. Each data point represents the average of technical replicates (n=3). Data are shown as mean ± SEM. **(E)** Genome browser tracks for MLL3 ChIP-Seq at the *CDKN2A* locus in HLE cells with sgGFP (control, black) or sgMLL3 (blue). **(F** and **G)** qPCR analysis for mRNA expression of **(F)** *INK4A* and **(G)** *ARF* in HLE cells with sgGFP (control, black) or sgMLL3 (blue). Each data point represents the average of technical replicates (n = 3). Data are shown as mean ± SEM.

Since the *Myc*; sgp53 and *Myc*; sgMll3 cells we studied above are not isogenic, we performed a series of additional experiments to demonstrate a direct transcriptional effect of MLL3 on the *Cdkn2a* locus. While p53 inactivation can lead to compensatory increases in *Ink4a* and *Arf* expression (Stott et al., 1998), *p53* suppression in sgMll3 cells produced only a subtle and inconsistent effect on expression of *Ink4a* and *Arf* (**Fig. S6A**), and *Mll3* suppression in sgp53 cells attenuated p16^Ink4a^ and p19^Arf^ protein levels (**Fig. S6B**). In addition, liver cancer cells produced by hydrodynamic delivery of the *Myc* transposon vector and *Axin1* sgRNAs (**Fig. S6C**) with or without an *Mll3* shRNA retained wild-type *p53* and showed reduced *Ink4a* and *Arf* mRNA and protein expression (**Fig. S6D-E**). These data imply that MLL3 supports a chromatin environment at the *Cdkn2a* locus that facilitates transcription of both *Ink4a* and *Arf* products of the *Cdkn2a* locus, and raises the possibility that these factors contribute to the tumor suppressor activity of MLL3 in liver cancer.

We next set out to determine whether MLL3 binding is sufficient to induce transcriptional activation of the *CDKN2A* locus and, in doing so, extend our analysis to human liver cancer cells. As the *MLL3* transcript is too large (14,733 bp) for cDNA transduction, we turned to the CRISPR activation (CRISPRa) system (Chavez et al., 2015) in a human hepatocellular carcinoma cell line (HLE). Following stable integration of the CRISPRa Cas9, cells were transduced with an sgRNA targeting the human *MLL3* promoter or sequences within GFP as a control (**Fig. 4C**). Cells expressing the *MLL3* sgRNA showed a marked and specific increase in the expression of endogenous *MLL3*, but not of *MLL4* or *TP53* (**Fig. 4D, Fig. S6F**), which was accompanied by a concomitant increase in MLL3 binding to the *CDKN2A* locus (**Fig. 4E)** and transcriptional upregulation of *INK4A* and *ARF* (**Fig. 4F-G**). Therefore, MLL3 directly binds and co-activates transcription of the *CDKN2A* locus in human liver cancer cells.

### MLL3 mediates oncogene-induced apoptosis in a *Cdkn2a*-dependent manner

The above results raise the possibility that the *Cdkn2a* products, *Ink4a* and *Arf*, may contribute to the tumor suppressive activity of MLL3. In this regard, *Myc* overexpression in primary cells (MEFs) often triggers apoptosis (Evan et al., 1992), and this in turn limits tumorigenesis in a manner that is dependent on *Cdkn2a* and most prominently *Arf* (Zindy et al., 1998). This pathway also suppresses liver tumorigenesis, since concomitant disruption of *Ink4a* and *Arf* using CRISPR, or through germline deletion of *Arf* alone, cooperated with *Myc* overexpression to rapidly promote tumor development (**Fig. S7A**). Similarly, *Mll3* suppression also attenuated MYC-induced apoptosis, as shown by tumor histology and apoptosis by TUNEL assay (Negoescu et al., 1997), 5 days after hydrodynamic delivery of transposon vectors encoding *Myc* together with GFP-linked shRNAs targeting *Mll3* (or Renilla luciferase as a control) (**Fig. 5A-B**). This dramatic difference in apoptosis correlated with an increase in retention of GFP-shMll3 expressing cells 10 days after injection (**Fig. S7B-C**). We also observed that apoptotic GFP-shRenilla-expressing hepatocytes were typically surrounded by immune cells, while shMll3-expressing cells formed incipiently transformed clusters that lacked immune infiltration (**Fig. S7D**). Altogether, these results show that *Mll3* suppression impairs *Myc*-induced apoptosis *in vivo* in a manner that is reminiscent of the anti-apoptotic effects of *Cdkn2a* loss in the context of aberrant *Myc* activation (Eischen et al., 1999; Jacobs et al., 1999; Schmitt et al., 1999).

**Figure 5.**
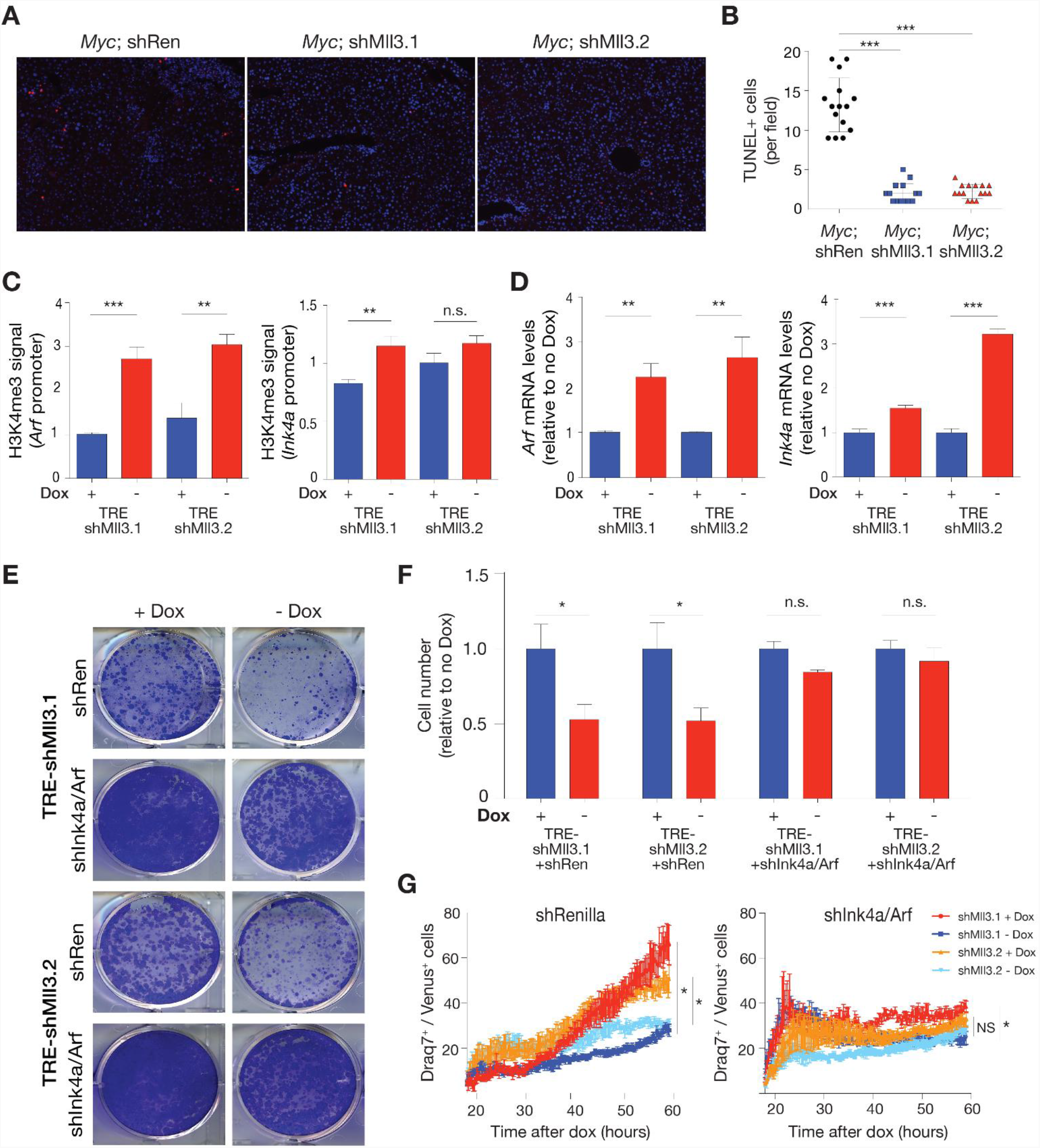
MLL3 mediates oncogene-induced apoptosis in a *Cdkn2a*-dependent manner. **(A)** Representative images of TUNEL-positive nuclei (red staining) in murine livers 5 days after hydrodynamic injection of indicated vector combinations. DAPI was used to visualize nuclei. **(B)** Quantification of TUNEL-positive nuclei in mouse livers 5 days after HTVI of indicated vector combinations. Data points represent the number of TUNEL-positive cells of 5 different high power fields in three independent murine livers per group. *** = *P* <0.001 (One-way ANOVA followed by post hoc t-tests). **(C)** ChIP-qPCR analysis for H3K4me3 signals at *Arf* and *Ink4a* promoters 4 days after dox withdrawal in *Myc*-rtTA3; TRE-shMll3 cells. Values are mean ± SD from technical replicates (n = 3) and the experiments were conducted in two independent LPC lines with different shMll3. **(D)** qPCR analysis for mRNA expression of *Arf* and *Ink4a* 4 days after dox withdrawal in two independent LPC lines with different shMll3. Values are mean ± SD from technical replicates (n = 3). *** = *P* < 0.001, ** = *P* < 0.01 (unpaired two-tailed t-test). **(E)** Representative images of colony formation assay of indicated cell lines 5 days after Dox withdrawal. **(F)** Quantification of colony formation assay. Values are mean ± SD of three independent experiments with two independent LPC lines. * = *P* < 0.05 (unpaired two-tailed t-test). **(G)** Time course analysis of Draq7 positive (viability dye) cells as a fraction of Venus positive, *Myc*-rtTA3; TRE-shMll3 cells expressing constitutive shRNAs targeting both *Ink4a/Arf* (shInk4a/Arf) or Renilla luciferase (shRen) off and on dox. Values represent mean ± SEM of triplicate wells of each genotype at each timepoint of two independently derived LPC lines, infected with either shRen or shInk4a/Arf. * = *P* < 0.05 (unpaired two-tailed t-test of final average percentage Draq7+/GFP+).

To model the interaction between *Myc* overexpression, *Mll3* function, and *Cdkn2a* regulation, we transduced liver progenitor cells (LPCs) with retroviral vectors encoding *Myc* linked to a reverse tetracycline transactivator (rtTA3), together with doxycycline (dox)-inducible *Mll3* shRNAs to enable reversible *Mll3* silencing (**Fig. S8A**). Infection of LPCs with *Myc* in the presence of *Mll3* (i.e. cells infected with *Myc*-rtTA3 and a dox-inducible shRNA targeting Renilla luciferase) acutely activated *Ink4a* and *Arf* expression (**Fig. S8B**) and these cells could not be maintained in culture. Phenocopying the ability of *Myc* and *Mll3* suppression to transform liver cells *in vivo*, combined *Myc* and shMll3 expression facilitated the persistent growth of cells maintained on Dox (**Fig. S8C-D**). By contrast, Dox withdrawal induced *Mll3* mRNA expression (**Fig. S8C**) and H3K4me3 deposition at the *Arf* and *Ink4a* promoters, ultimately leading to elevations in *Arf* and *Ink4a* mRNA and protein (**Fig. 5C-D, Fig. S8F**), reduced colony formation, and increased apoptosis (**Fig. S8D-E**). Furthermore, constitutive shRNA-mediated knockdown of *Arf* and *Ink4a* through targeting of the shared exon 2 (shInk4a/Arf) significantly rescued colony forming capacity and prevented cell death following *Mll3* restoration as determined by time-lapse microscopy of cells cultured with a fluorescent viability dye (**Fig. 5E-G, Fig. S8G**). These data support a model whereby a prominent tumor suppressive output of MLL3 in liver cancer involves direct upregulation of *Cdkn2a* that, when impaired, attenuates the MYC-induced apoptotic program and permits tumor progression.

## DISCUSSION

Our study combined genetic, epigenomic, and animal modeling approaches to identify *Cdkn2a* as an important regulatory target of MLL3 in both mouse and human liver cancers. Our results support a model whereby oncogenic stress, herein produced by MYC, leads to an increase in the binding of MLL3 to the *CDKN2A* locus, an event that is associated with the accumulation of histone marks linked to the biochemical activity of MLL3-containing complexes and conducive to gene activation (**Fig. 6**). Accordingly, these events are accompanied by transcriptional upregulation of two key *CDKN2A* gene products, *INK4A* and *ARF*, and suppression of *MLL3* phenocopies the effects of *CDKN2A* inactivation in abrogating MYC-induced apoptosis. Conversely, suppression of *CDKN2A* abolishes the anti-proliferative effects of *MLL3* restoration. As such, our results establish a conserved epistatic relationship between the chromatin modifier MLL3 and a well-characterized tumor suppressor network.

**Figure 6.**
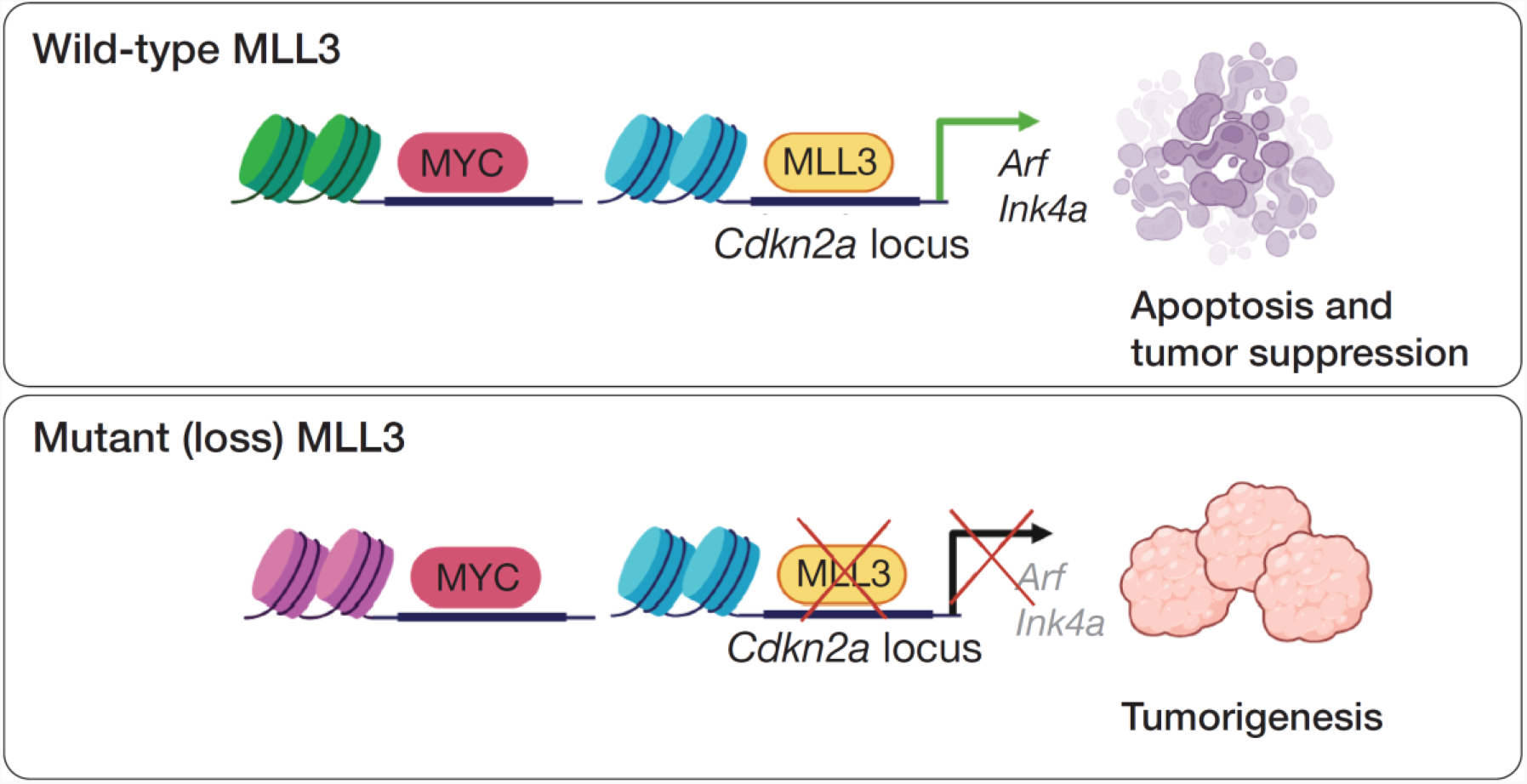
Model of MLL3 as a tumor suppressor in liver cancer. MLL3 acts as a tumor suppressor to restrict MYC-induced liver tumorigenesis by directly activating the *Cdkn2a* locus to mediate tumor cell apoptosis.

While the epistatic relationship described above is supported by the trends towards mutually exclusivity of *MLL3* and *CDKN2A* alterations in liver cancer and several other tumor types, this was not always the case. In some tumor types, this may be due to limited samples sizes, other functionally important components linked to the *CDKN2A* locus could produce *CDKN2A*-independent forces that drive selection for chromosome 9p deletions, including type I interferon genes, *CDKN2B*, and *MTAP* (Fountain et al., 1992; Schmid et al., 2000; Xia et al., 2021). Alternatively, mutual exclusivity between *MLL3* and *CDKN2A* alterations would only be expected under circumstances where *CDKN2A* action is the most dominant MLL3 effector, and it seems likely that multiple downstream genes contribute to tumor suppression and their relative importance may vary between cell and tissue types. Such a variable output in cancer relevant gene regulation has been noted for other chromatin regulators that, at the extreme, serve as pro-oncogenic factors in some contexts and tumor suppressors in others (Ntziachristos et al., 2012; Soto-Feliciano et al., 2017; Souroullas et al., 2016; Tirode et al., 2014).

UTX (KDM6A), MLL3 (KMT2C), and MLL4 (KMT2D), the core catalytic components of the COMPASS-like complex, are all considered tumor suppressors with frequent loss-of-function genomic alterations found in a broad spectrum of human cancers (Sze and Shilatifard, 2016); (Revia et al., 2021). While each of these components regulate redundant sets of genes (Hu et al., 2013; Lee et al., 2009), it appears that they may exert their tumor suppressive functions through different mechanisms. In liver and pancreas cancer models, UTX can control the expression of negative regulators of mTOR such as *DEPTOR*, and its disruption prevents their transcription and facilitates tumorigenesis through increased mTORC1 activity (Revia et al., 2021). Additionally, while the mechanisms of MLL4 activity in liver cancer have not been examined, studies suggest that MLL4 suppresses skin carcinogenesis by promoting lineage stability and ferroptosis independently of MLL3 (Egolf et al., 2021). Our study demonstrates that MLL3 is both necessary and sufficient for efficient transcriptional activation of the *CDKN2A* locus that drives oncogene-induced apoptosis. The molecular basis for this heterogeneity in effector output remains to be determined, but it seems likely that different subsets of target genes are preferentially disabled by haploinsufficiency of individual components and/or subject to compensation by remaining COMPASS complex activities. Systematic studies comparing the binding, histone modifications, and transcriptional output of cells across a spectrum of allelic configurations of COMPASS complex factors will be needed to achieve a more holistic understanding of their functions and interactions in different contexts.

The most well-established role for MLL3/4-UTX-containing complexes is the control of H3K4 mono-methylation at enhancers during development (Herz et al. 2010; Hu et al. 2013). While our ChIP-Seq studies also revealed binding of MLL3/4 to enhancers in liver tumor cells, an even larger fraction of genes – including *Cdkn2a* – showed MLL3/4 chromatin enrichment at gene promoters and, indeed, transcription of this class of genes was most affected by *Mll3* disruption. Interestingly, *Mll3* suppression preferentially limited the MLL3/4 enrichment at promoters and shifted residual complex binding towards intergenic regions. Such dynamic regulation of distinct cis-acting elements by the MLL3/4 complex has also been observed in other contexts (Cheng et al., 2014; Soto-Feliciano et al., 2021), where the non-canonical binding of MLL3/4 at promoters is a recurrent tumor suppressive mechanism in cancer cells. Further studies into the action and regulation of MLL3/4 complexes at promoters will be informative and may shed new insights into the actions of the COMPASS-like complex in cancer.

While the dominant role of *CDKN2A* in mediating the tumor suppressive effects of the broadly acting MLL3 enzyme is surprising, the contribution of a single gene to the functional output of chromatin-complex disruption is not unprecedented. Indeed, Polycomb Repressive Complexes (PRC) broadly repress gene expression in different cell types through the coordinated action of PRC1 and PRC2 complexes that deposit and maintain repressive H3K27me3 marks on the enhancers of target genes, including *CDKN2A (Bracken et al*., *2007; Kotake et al*., *2007)*. Despite these similarly broad effects, *CDKN2A* is often the most functionally relevant target of PRC-mediated repression, as genetic deletion of either the PRC1 component *Bmi1* or the PRC2 component *Ezh2*, or treatment with small molecule inhibitors of EZH2, can facilitate *Cdkn2a* induction in normal and tumor cells. This, in turn, triggers anti-proliferative responses that can be rescued by *Cdkn2a* deletion (Jacobs et al., 1999; Richly et al., 2011). It is noteworthy that the COMPASS-like complexes are biochemically and functionally similar to *Trithorax* complexes in *Drosophila*, which have an evolutionarily conserved antagonistic relationship with Polycomb Repressive Complexes (PRC1 and PRC2) that controls epigenetic memory and cell fate during development (Mills, 2010; Piunti and Shilatifard, 2016). Our findings suggest such antagonism extends to tumor suppression in mammalian cells, likely via regulation of *Cdkn2a* and other tumor suppressor genes (Soto-Feliciano et al., 2021).

## ACKNOWLEDGEMENTS

We thank Charles Sherr for constructive guidance and advice on all aspects of this study. We thank Ali Shilatifard, Lu Wang, and all members of the Lowe lab for helpful and stimulating discussions. We gratefully thank A. Chramiec for excellent technical assistance. We thank Joanna Wysocka (Stanford University) for kindly sharing the anti-MLL3/4 antibody used in our ChIP-Seq experiments. This work was supported by grants to S.W.L. (P01 CA013106 and R01 CA233944) from the NIH/NCI, as well as by the National Center for Tumor Disease, Heidelberg, and grants of the German Research Foundation to D.F.G. (SFB/TRR77). This work was also supported by the NIH/NCI Cancer Center Support Grant to Memorial Sloan Kettering Cancer Center (P30 CA008748). Y.M.S.F. is supported by a MOSAIC K99/R00 Award from the NIH/NIGMS (1K99GM140265-01). C.Z. is supported by an F32 Postdoctoral Fellowship (1F32CA257103) from the NIH/NCI. J.P.M. was a recipient of a Postdoctoral Fellowship (PF-14-066-01-TBE) from the American Cancer Society. D.F.T. is supported by a Young Investigator Group (VH-NG-1114) by the Helmholtz foundation. S.W.L. is the Geoffrey Beene Chair for Cancer Biology and an investigator of the Howard Hughes Medical Institute.

## AUTHOR CONTRIBUTIONS

D.F.T., J.P.M.IV, C.H.H., and S.W.L. conceived the project. D.F.T., Y.M.S.F., C.Z., J.P.M.IV, C.H.H., and A.B. designed and analyzed experiments. D.F.T., Y.M.S.F., C.Z., J.P.M.IV, C.H.H., A.B., A.S., and S.T. performed experiments. C.W.C. and G.L. contributed important and unpublished reagents to the study. Y.H. and C.C.C. analyzed RNA-Seq data. Y.M.S.F., R.P.K., C.W.C. and S.A.A. analyzed and supervised ChIP-Seq experiments. Y.H. and C.Z. analyzed human data. C.D.A provided conceptual advice. Y.M.S.F., C.Z., D.F.T., C.H.H., J.P.M.IV, and S.W.L. wrote the paper with input from all co-authors.

## COMPETING INTERESTS

S.W.L. is an advisor for and has equity in the following biotechnology companies: ORIC Pharmaceuticals, Faeth Therapeutics, Blueprint Medicines, Geras Bio, Mirimus Inc., and PMV Pharmaceuticals. S.W.L. also acknowledges receiving funding and research support from Agilent Technologies and Calico, for the purposes of massively parallel oligo synthesis and single-cell analytics, respectively. C.D.A. is a co-founder of Chroma Therapeutics and Constellation Pharmaceuticals and a Scientific Advisory Board member of EpiCypher. The remaining authors declare no competing interests.

## MATERIALS AND METHODS

### Animal experiments

8-10 weeks old female C57BL/6 animals were purchased from Envigo (formerly Harlan). *Arf*-null (p19^-/-^) animals (C57BL/6 background), originally provided by Dr. Charles Sherr, St. Jude Children’s Research Hospital, were maintained in our breeding colony. For hydrodynamic tail vein injection, a sterile 0.9% NaCl solution/plasmid mix was prepared containing 5 µg DNA of pT3-*Myc* and either 20 µg of pX330 expressing indicated sgRNAs or 20 µg of pT3-EF1a-GFP-miRE plasmid together with CMV-SB13 Transposase (1:5 ratio). Mice were randomly assigned to experimental groups and injected with the 0.9% NaCl solution/plasmid mix into the lateral tail vein with a total volume corresponding to 10% of body weight in 5-7 seconds as described before ((Largaespada, 2009; Moon et al., 2019; Tschaharganeh et al., 2014; Xue et al., 2014)). Injected mice were monitored for tumor formation by abdominal palpation. All animal experiments were approved by the MSKCC Institutional Animal Care and Use Committee (protocol 11-06-011).

### Vector constructs

The pT3-*Myc* vector and pT3-EF1a-GFP-miRE plasmid were described before (Huang et al., 2014). For CRISPR/Cas9-mediated genome editing, sgRNAs were subcloned into pX330 (Addgene, #42230) (Hsu et al., 2013). All shRNA and sgRNA sequences are listed in **Supplementary Table S1**.

### Derivation of primary liver tumor cell lines

Liver tumors were resected with sterile instruments, and 10-50 mg of tumor tissue was minced and washed in sterile PBS, incubated in a mix of 4 mg/mL collagenase IV and dispase (w/v in sterile, serum free DMEM) with gentle shaking, washed with PBS, incubated for 5 min in 0.05% trypsin, and washed and plated in complete DMEM (10%FBS, 1x Penicillin/Streptomycin) on collagen coated plates (PurCol, Advanced Biomatrix). Primary cultures were passaged until visibly free from fibroblasts.

### Analysis of CRISPR-directed mutations

CRISPR mediated insertions and deletions were detected by Surveyor assay as directed by the manufacturer (Transgenomic/IDT). Briefly, genomic DNA was extracted from primary tumors and cell lines by isopropanol precipitation following overnight lysis at 37°C in buffer containing 0.4 mg/ml Proteinase K, 10mM Tris, 100mM NaCl, 10mM EDTA, and 0.5% SDS, pH 8.0. ∼250-500 bp regions flanking predicted CRISPR cleavage sites were PCR amplified with Herculase II taq polymerase, column purified (Qiagen), heated to 95°C, and slowly cooled to promote annealing of heteroduplexes. Following treatment with Surveyor nuclease, products were analyzed by electrophoresis on a 2% polyacrylamide gel. Primers used for Surveyor Assay are listed in Supplementary Table S1. Amplified PCR products were separately gel purified and ligated into blunt-end digested pBlueScript (Stratagene). Sanger nucleotide sequencing analysis was performed on DNA from 48 transformed colonies using a T7 primer.

### CRISPR activation (CRISPRa)

Human HCC cell line HLE was purchased from JCRB Cell Bank (JCRB0404), which was transduced by the lentivirus expressing nuclease-dead Cas9 (dCas9) fused with VP64-p65-Rta (VPR) (Chavez et al., 2015) and sgRNA against *MLL3/KMT2C* (sequence in Supplementary Table S1) to generate stable *MLL3* CRISPRA HLE line.

### Generation and modification of primary cells

Liver progenitor cells (LPCs) from E13.5-15.5 C57BL/6 embryos were isolated and grown in hepatocyte growth media (HGM) as previously described (Zender et al., 2005). To simultaneously overexpress Myc and conditionally suppress Mll3, LPCs were co-infected with a retroviral construct constitutively expressing both Myc and a reverse tet-transactivator (rtTA) (MSCV-Myc-IRES-rtTA) along with retroviral TRMPV vectors (MSCV-TRE-dsRed-miR30/shRNA-PGK-Venus-IRES-NeoR) (Zuber et al., 2011) expressing ds-Red linked, tet-responsive shRNAs targeting Mll3 cloned into an optimized mir-30 context (“mir-E”, TRPMVe) (Fellmann et al., 2013). For selection of infected cells and sustained shMll3 expression, cells were maintained in HGM with neomycin (1 mg/mL) and doxycycline (1 ug/mL) starting 2 days after infection. To introduce constitutively expressed shRNAs in the setting of inducible shMll3, retroviral MLPe vectors (MSCV-LTR-mir-E-PGK-Puro-IRES-GFP) (Dickins et al., 2005), GFP-linked shRNAs targeting either *Cdkn2a* or a control shRNA targeting Renilla luciferase were co-infected with MSCV-Myc-ires-rtTa and TRMPVe-shMLL3. Triple infected cells were maintained in media with neomycin, puromycin (2 µg/mL), and doxycycline 2 days post infection. Infected continuously proliferating cells were transitioned to growth in complete DMEM, and maintained on collagen-coated plates.

### Colony assays

For measurement of cell proliferation, 5000 transduced and selected LPCs or MEFs were plated in triplicate in 6 well plates. Tetracycline inducible-shMLL3 expressing LPCs were grown in the presence or absence of doxycycline, and cells were fixed with formalin and methanol and stained with 0.05% crystal violet after 5 days. 5000 MEFs were plated in 6-well plates, fixed after 6 days with formalin and methanol, and stained with 0.05% crystal violet.

### Apoptosis assays

Apoptosis was measured in LPCs via Annexin V staining according to manufacturer’s instructions (eBiosciences, Annexin-V APC). 25,000 cells were grown with and without doxycycline for 3 days, trypsinized, washed with Annexin-V binding buffer, and ∼100,000 cells were incubated with Annexin-V APC and analyzed on an LSRII flow cytometer (BD).

### Live Imaging

1000 LPCs immortalized by linked overexpression of myc and 2 independent, inducible Mll3 shRNAs constitutively expressing shRNAs targeting Renilla luciferase or Cdkn2a (generated as detailed above) were plated on collagen coated, 96 well, clear bottom imaging plates in media supplemented with 300nM Draq7 (Invitrogen) with and without doxycycline, in triplicate by genotype. 18 hours after plating cells, Venus (marking all plated cells) and Draq7 fluorescence was collected in 2, 10X fields of each well every 15 minutes for 41 hours using an automated, high content microscope (InCell 6000, General Electric).

### Chromatin immunoprecipitation (ChIP)

Histone ChIP was performed as previously described (Lee et al., 2006). Briefly, cell samples were cross-linked in 1% formaldehyde for 10 minutes, and the reaction was stopped by addition of glycine to 125 mM final concentration. Fixed cells were lysed in SDS lysis buffer, and the chromatin was fragmented by sonication (Covaris). Sheared chromatin was incubated with a final 10 μg/mL concentration of antibodies against either H3K4me3 (Abcam, ab8580, Lot:GR164706-1), H3K27ac (Abcam, ab4729, Lot:GR200563-1), H3K4me1 (Abcam; ab8895, Lot:GR114265-2) or normal rabbit IgG (Abcam, ab46540) at 4 °C for overnight. Antibodies were recovered by binding to protein A/G agarose (Millipore), and the eluted DNA fragments were used directly for qPCR or subjected to high-throughput sequencing (ChIP-Seq) using a HiSeq 2000 platform (Illumina). High-throughput reads were aligned to mouse genome assembly NCBI37/mm9 as previously described (Barradas et al., 2009). Reads that aligned to multiple loci in the mouse genome were discarded. The ChIP-Seq signal for each gene was quantified as total number of reads per million (RPM) in the region 2 kb upstream to 2 kb downstream of the transcription start site (TSS). The complete dataset is available at NCBI Gene Expression Omnibus (GSE85055). Primers used for ChIP-qPCR of mouse *Cdkn2a* promoter (Barradas et al., 2009) are included in the Supplementary Table S1.

For the MLL3 ChIP-Seq, the following protocol was used. Cross-linking ChIP in mouse and human HCC cells was performed with 10-20×10^7^ cells per immunoprecipitation. Cells were collected, washed once with ice-cold PBS, and flash-frozen. Cells were resuspended in ice-cold PBS and cross-linked using 1% paraformaldehyde (PFA) (Electron Microscopy Sciences) for 5 minutes at room temperature with gentle rotation. Unreacted PFA was quenched with glycine (final concentration 125mM) for 5 minutes at room temperature with gentle rotation. Cells were washed once with ice-cold PBS and pelleted by centrifugation (800g for 5 minutes). To obtain a soluble chromatin extract, cells were resuspended in 1mL of LB1 (50mM HEPES pH 7.5, 140mM NaCl, 1mM EDTA, 10% glycerol, 0.5% NP-40, 0.25% Triton X-100, 1X complete protease inhibitor cocktail) and incubated at 4°C for 10 minutes while rotating. Samples were centrifuged (1400g for 5 minutes), resuspended in 1mL of LB2 (10mM Tris-HCl pH 8.0, 200mM NaCl, 1mM EDTA, 0.5mM EGTA, 1X complete protease inhibitor cocktail), and incubated at 4°C for 10 minutes while rotating. Finally, samples were centrifuged (1400g for 5 minutes) and resuspended in 1mL of LB3 (10mM Tris-HCl pH 8.0, 100mM NaCl, 1mM EDTA, 0.5mM EGTA, 0.1% sodium deoxycholate, 0.5% N-Lauroylsarcosine, 1X complete protease inhibitor cocktail). Samples were homogenized by passing 7-8 times through a 28-gauge needle and Triton X-100 was added to a final concentration of 1%. Chromatin extracts were sonicated for 14 minutes using a Covaris E220 focused ultrasonicator. Lysates were centrifuged at maximum speed for 10 minutes at 4°C and 5% of supernatant was saved as input DNA. Beads were prepared by incubating them in 0.5% BSA in PBS and antibodies overnight (100μL of Dynabeads Protein A or Protein G (Invitrogen) plus 20μL of antibody). Antibody used the anti-MLL3/4 (kindly provided by the Wysocka laboratory (Dorighi et al., 2017)). Antibody-Beads mixes were washed with 0.5% BSA in PBS and then added to the lysates overnight while rotating at 4°C. Beads were then washed six times with RIPA buffer (50mM HEPES pH 7.5, 500mM LiCl, 1mM EDTA, 0.7% sodium-deoxycholate, 1% NP-40) and once with TE-NaCl Buffer (10mM Tris-HCl pH 8.0, 50mM NaCl, 1mM EDTA). Chromatin was eluted from beads in Elution buffer (50mM Tris-HCl pH 8.0, 10mM EDTA, 1% SDS) by incubating at 65°C for 30 minutes while shaking, supernatant was removed by centrifugation, and crosslinking was reversed by further incubating chromatin overnight at 65°C. The eluted chromatin was then treated with RNaseA (10mg/mL) for 1 hour at 37°C and with Proteinase K (Roche) for 2 hours at 55°C. DNA was purified by using phenol-chloroform extraction followed with ethanol precipitation. The NEBNext Ultra II DNA Library Prep kit was used to prepare samples for sequencing on an Illumina NextSeq500 (75bp read length, single-end, or 37bp read length, paired-end).

### Immunoblotting

Cell pellets were lysed in Laemmli buffer (100 mM Tris-HCl pH 6.8, 5% glycerol, 2% SDS, 5% 2-mercaptoethanol). Equal amounts of protein were separated on 12% SDS–polyacrylamide gels and transferred to PVDF membranes (90V, 75 mins). β-actin was used as a control to ensure equal loading, and images were analyzed using the AlphaView software (ProteinSimple). Immunoblotting was performed using antibodies for MYC (1:1000, Abcam, ab32072), p53 (1:500, Leica Biosystems, NCL-p53-505), p19 (1:250, Santa Cruz Biotechnology, sc-32748), p16 (1:250, Santa Cruz Biotechnology, sc-1207), Axin1 (1:1000, Cell Signaling, #2074), β-actin (1:10000, Sigma-Aldrich, clone AC-15).

### Quantitative RT-PCR

Total RNA was isolated using RNeasy Mini Kit, QIAshredder Columns and RNase-Free DNase Set (Qiagen). cDNA synthesis was performed using TaqMan® Reverse Transcription Reagents (Thermo Fisher Scientific). Real-time PCR was carried out using Power SYBR® Green Master Mix (Thermo Fisher Scientific) and the Life Technologies ViiA™ 7 machine. Transcript levels were normalized to the levels of mouse or human *Actb* mRNA expression and calculated using the ΔΔCt method. Each qRT-PCR was performed in triplicate using gene-specific primers (sequences listed in Supplementary Table S1).

### RNA sequencing and differential expression analysis

For RNA sequencing, total RNA from three independent tumor-derived cell lines (*Myc*; sgp53 and *Myc*; sgMll3) was isolated using RNeasy Mini Kit, QIAshredder Columns and RNase-Free DNase Set (Qiagen). RNA-Seq library construction and sequencing were performed according to protocols used by the integrated genomics operation (IGO) Core at MSKCC. 5-10 million reads were acquired per replicate sample. After removing adaptor sequences with Trimmomatic, RNA-seq reads were aligned to GRCm38.91(mm10) with STAR (Dobin et al., 2013). Genome wide transcript counting was performed by HTSeq to generate a FPKM matrix (Anders et al., 2015). Differentially expressed genes were identified by DESeq2 (v.1.8.2, package in R) and plotted in the volcano plot. The complete dataset is available at NCBI Gene Expression Omnibus (GSE85055).

### Human cancer analyses

RNA sequencing data of selected samples with somatic mutations and homozygous deletions of either *MLL3*/*KMT2C, CDKN2A, TP53*, and *RB1* in the TCGA hepatocellular carcinoma (HCC) dataset were downloaded from Broad Institute TCGA Genome Data Analysis Center. To obtain transcriptional signatures of HCC with genomic mutations and deletions of either *MLL3*/*KMT2C, CDKN2A*, and *RB1*, differential gene expression analyses were performed by DESeq2 (with *TP53*-mutated HCCs as controls). The Oncoprints of homozygous deletions and somatic mutations of *MLL3*/*KMT2C, CDKN2A*, and *TP53*, as well as MYC gains and amplifications from human HCC datasets [TCGA ((Schulze et al., 2015; The Cancer Genome Atlas Research Network, David A. Wheeler, Lewis R. Roberts, 2017)), MSK, (Schulze et al., 2015), and RIKEN (Fujimoto et al., 2012)] and all other cancer types were generated by cBioPortal (www.cbioportal.org) (Cerami et al., 2012; Gao et al., 2013).

### Gene set enrichment analysis (GSEA) analyses

Gene set enrichment analysis (GSEA) was performed using the GSEAPreranked tool for conducting gene set enrichment analysis of data derived from RNA-seq experiments (version 2.07) against other signatures. The metric scores (normalized enrichment scores and false discovery rate q-values) were calculated using the sign of the fold change multiplied by the inverse of the *P*-value (Subramanian et al., 2005). Specifically, transcriptional signatures of human HCCs with genomic inactivation of *CDKN2A* were compared to other human HCCs with *MLL3* or *RB1* alterations, and mouse HCC cell lines (*Myc*; sgMll3 vs *Myc*; sgp53, genes with log_2_(fold-change)>4).

### Statistical analysis

Data are presented as mean ± standard deviation (SD) or standard error of the mean (SEM) as specified. The Spearman rank coefficient was used for statistical measure of association as indicated. The statistical comparison between 2 groups was accomplished with the two-tailed student’s t-test, or One-way ANOVA followed by post hoc t-tests among 3 or more groups. The analyses for co-occurrence or mutual exclusivity were performed using Fisher Exact Test. All statistical tests were performed using the Prism 8 software. The investigators were not blinded to the groups for the experiments. No samples or animals were excluded from the analysis.

