## Supplementary figures and table for "MLL3 regulates the *CDKN2A* tumor suppressor locus in liver cancer"

**Supplementary Figure 1. Suppression of *Mll3* by CRISPR or RNAi promotes *Myc*-driven liver cancer.**

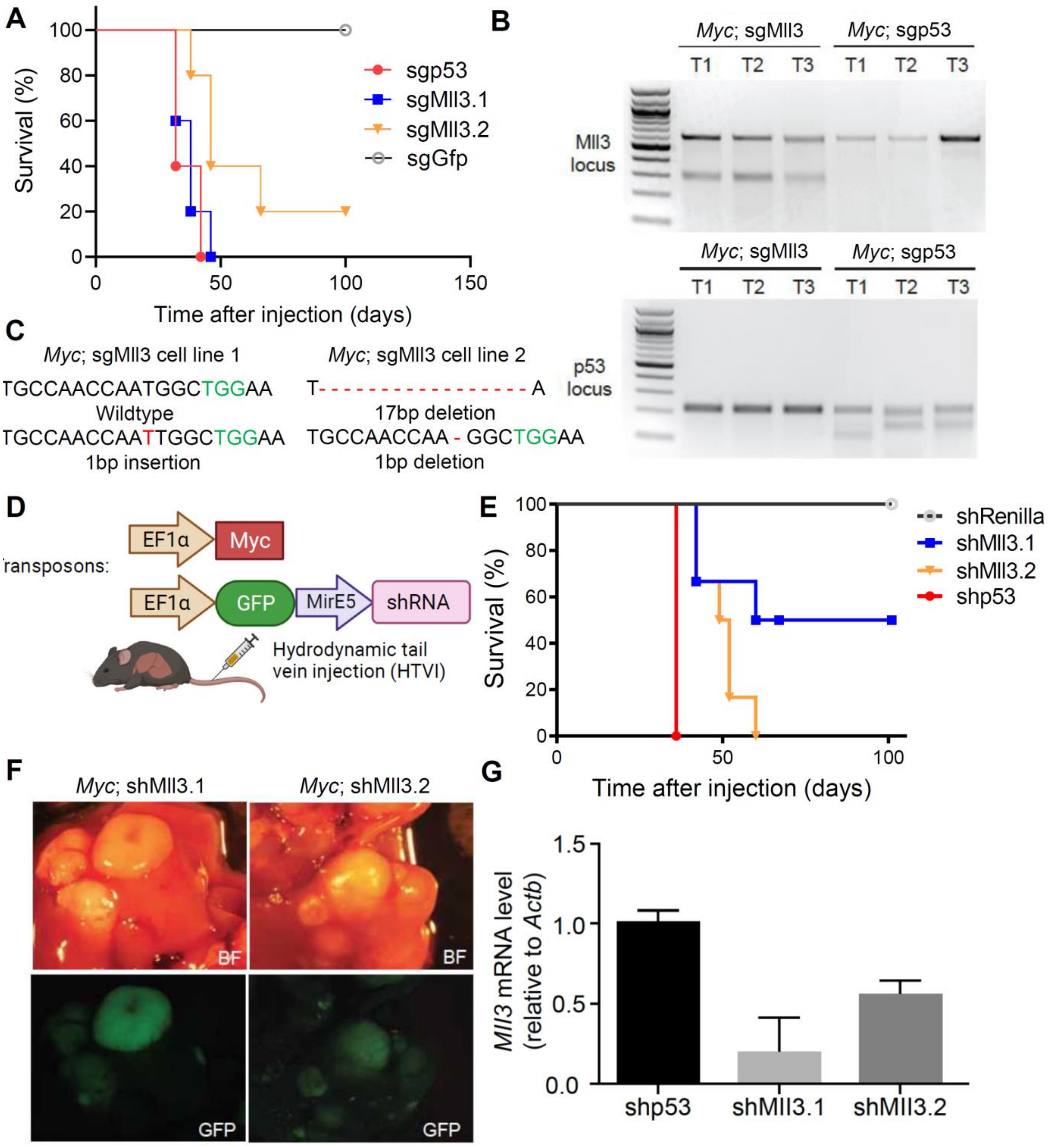


**(A)** Survival curve of a second cohort of mice injected with *Myc* transposon and pX330 expressing two independent sgRNAs targeting *Mll3* (*Myc;* sgMll3.1, n = 5; *Myc;* sgMll3.2, n = 5). *Myc;* sgp53 (n = 5), and

*Myc;* sgGFP (n = 5) serve as controls. **(B)** Surveyor assay of PCR amplicons flanking predicted *Mll3* and *p53* CRISPR cleavage sites in *Myc;* sgMll3 and *Myc;* sgp53 cells. The ratio of intensity of the lower bands (containing indels) to that of the upper band (untargeted) provides an estimate of the efficiency of CRISPR-Cas9 cleavage at the *Mll3* and *Tpr53* loci. **(C)** Sanger sequencing of the CRISPR targeted *Mll3* locus from two independently derived cell lines using two different sgMll3. The PAM-sequence is indicated in green, indels are indicated in red. **(D)** Schematic for hydrodynamic tail vein injection (HTVI) for in vivo gene silencing in mouse livers. HTVI was used to introduce transposons enforcing stable expression of Myc and GFP-linked, constitutive shRNAs targeting genes of interest. **(E)** Survival curve of mice injected with transposon vectors expressing Myc oncogene and two independent shRNAs targeting Mll3 (*Myc;* shMll3.1, n = 6; *Myc;* shMll3.2, n = 6), p53 (*Myc;* shp53, n = 3), and Renilla luciferase (*Myc*; shRen, n = 3). **(F)** Representative images of tumor nodules observed in *Myc;* shMll3 mice expressing shRNA-linked GFP. **(G)** *Mll3* mRNA expression measured by qPCR in indicated tumor tissues. Values are mean ± SD from experimental triplicates in 3 different tumors of each group.

**Supplementary Figure 2. *Mll3* deficiency disrupts its binding at promoters in liver cancer cells.**

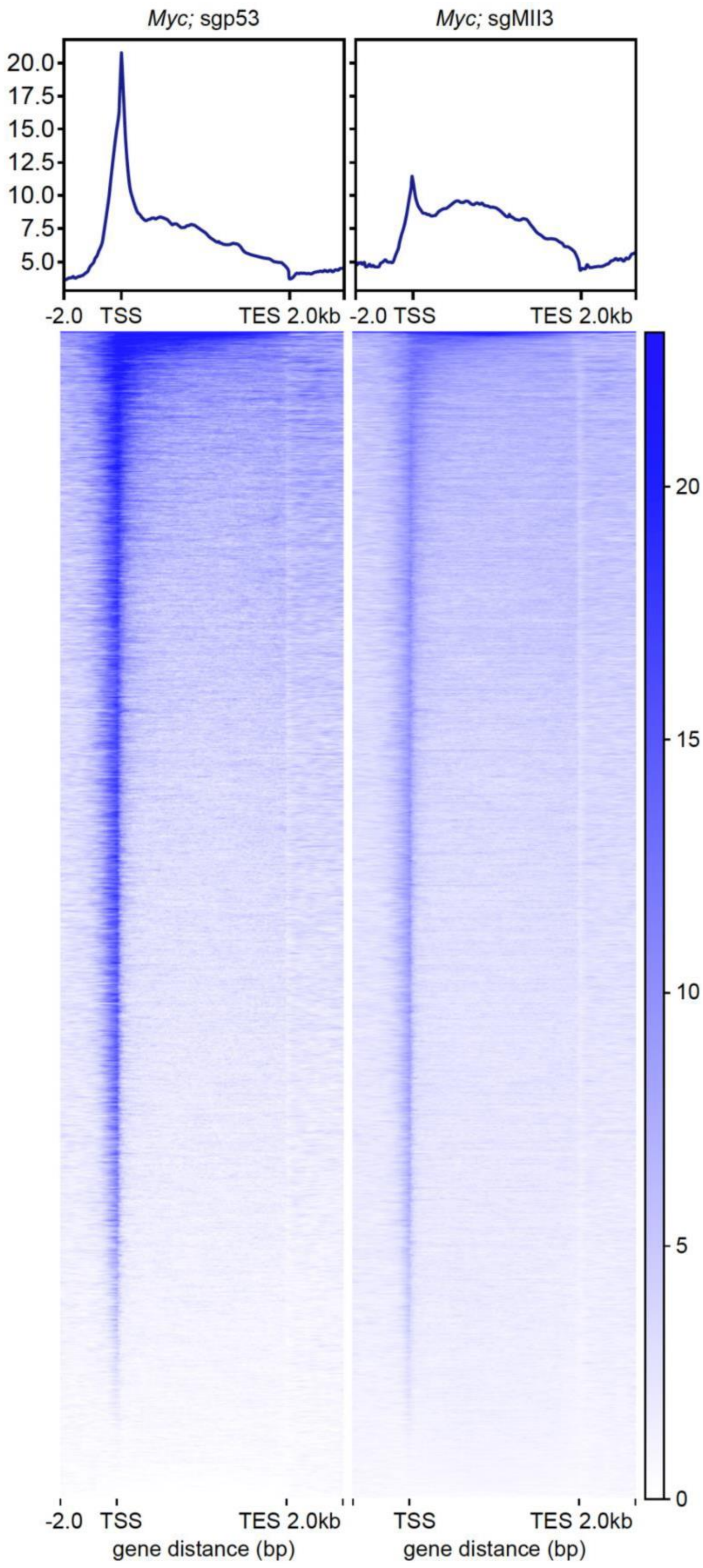


Tornado plots showing MLL3 ChIP-Seq signals at promoter regions in *Myc*; sgp53 and *Myc*; sgMll3 cells. Promoter regions were defined as transcriptional start sites (TSS) ± 2kb. TES, transcription end sites.

**Supplementary Figure 3.** ***Mll3*** **disruption impacts transcriptional and histone modification profiles in liver tumors.**

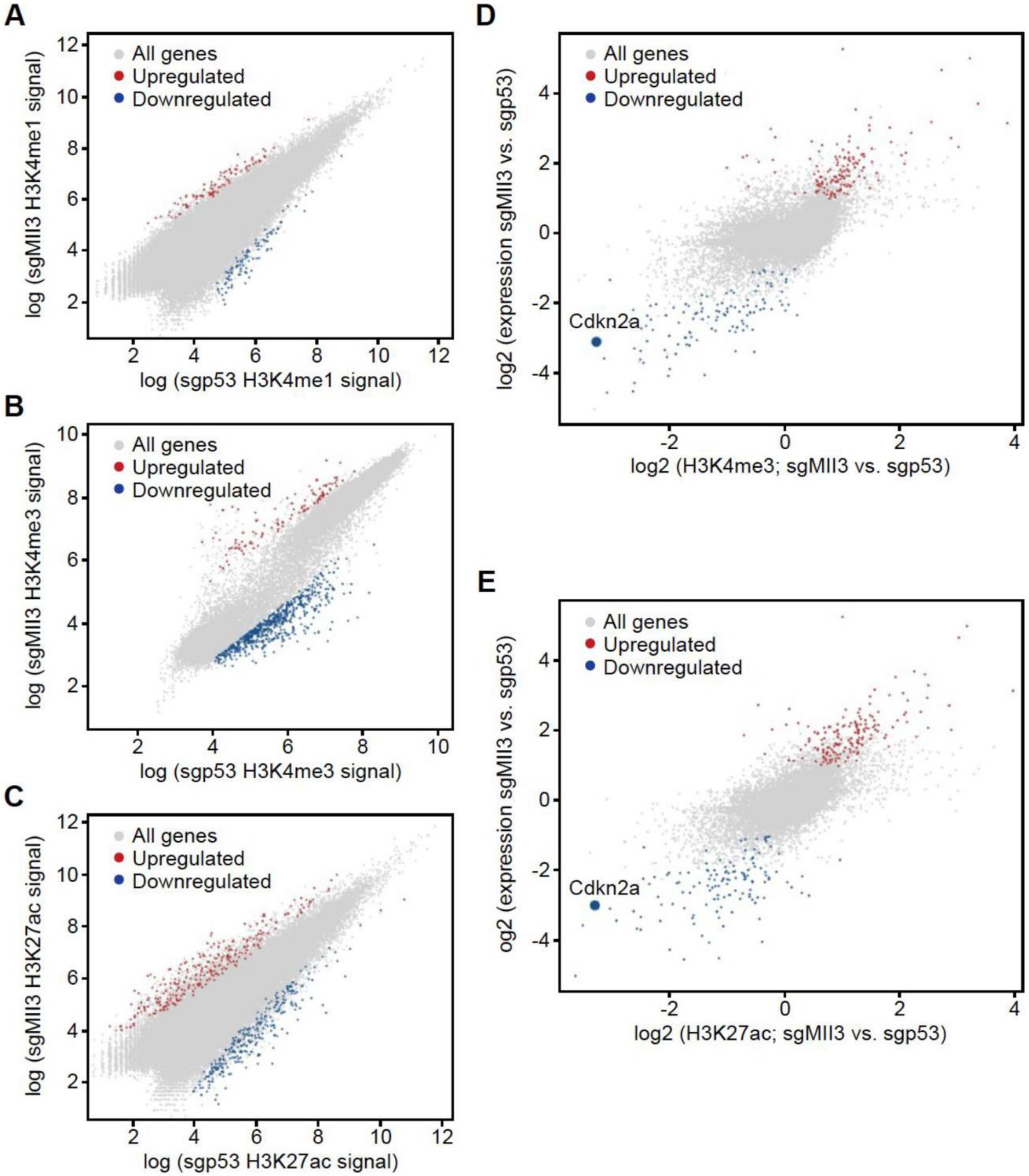


**(A-C)** Scatter plots comparing ChIP-Seq signals for **(A)** H3K4me1, **(B)** H3K4me3, and **(C)** H3K27ac in *Myc;* sgMll3 (y-axis) and *Myc;* sgp53 (x-axis) liver tumor-derived cell lines. **(D** and **E)** Scatter plots comparing RNA-seq (y-axis) and ChIP-Seq signals (x-axis) for **(D)** H3K4me3 and **(E)** H3K27ac in *Myc;* sgMll3 HCC cell lines.

**Supplementary Figure 4. The relationship between the genetic alterations of *MLL3* and *CDKN2A* in human cancers.**

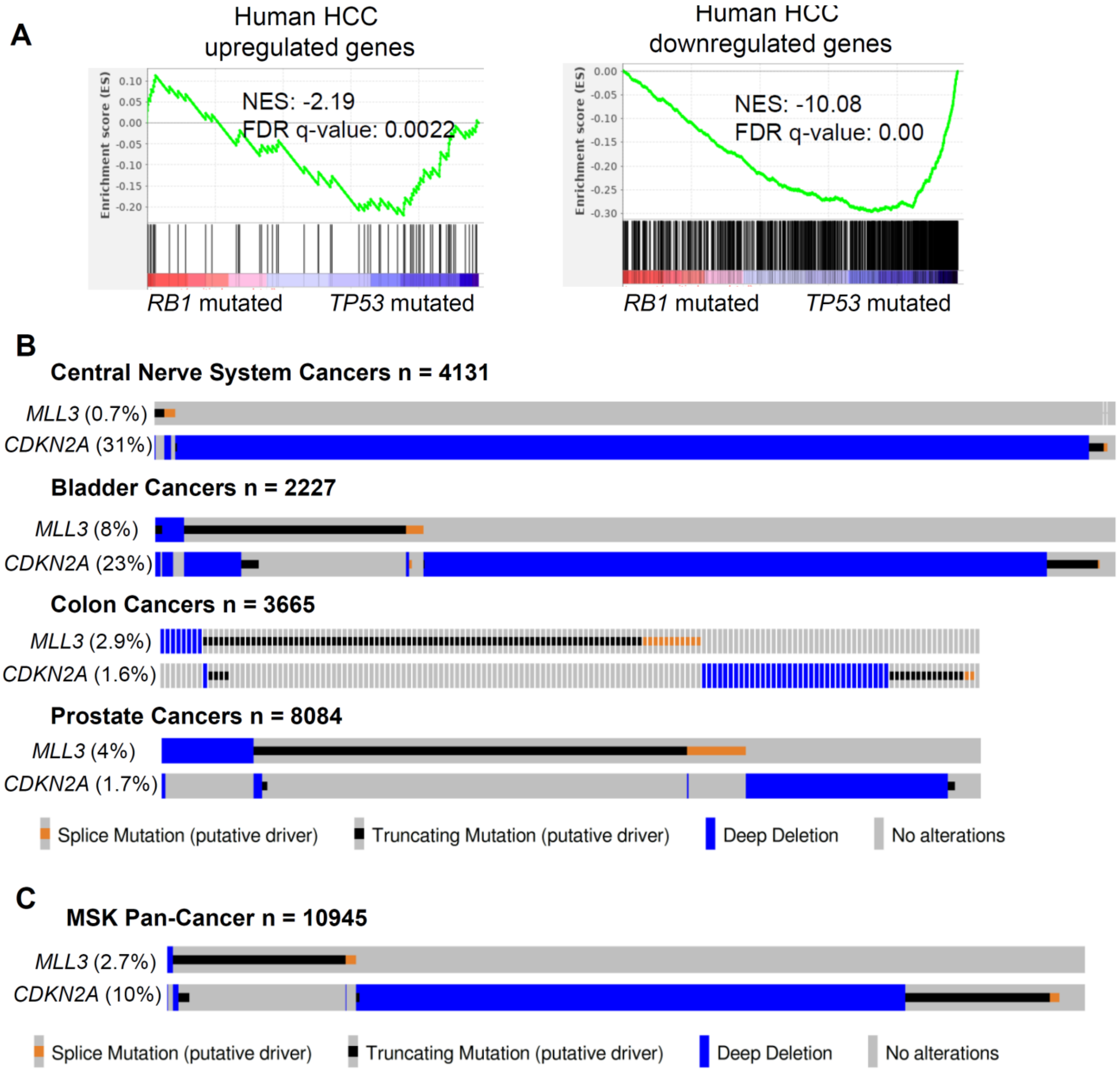

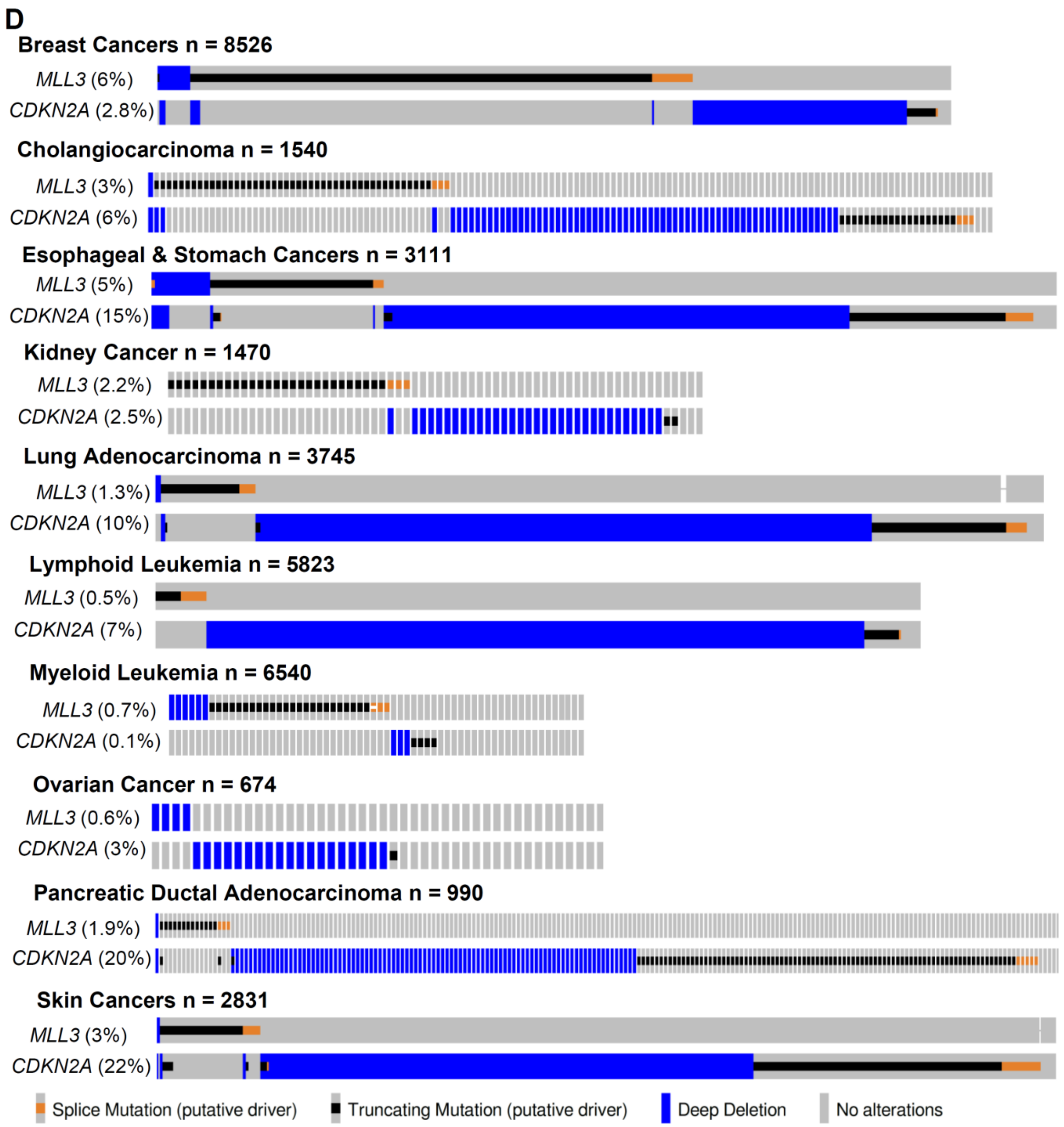


**(A)** Gene set enrichment analysis (GSEA) plots of human HCC transcriptional signatures with *CDKN2A* mutations or homozygous deletions in human HCCs with mutations and deletions of *RB1*. Normalized enrichment scores (NES) and False discovery rate (FDR) q-values were calculated by GSEA. **(B)** Oncoprints displaying the co-occurrence between the genomic alterations (truncating mutations and deep deletions) of *MLL3* and *CDKN2A* in four human cancer types. **(C** and **D)** Oncoprints displaying the mutual exclusivity between the genomic alterations (truncating mutations and deep deletions) of *MLL3* and *CDKN2A* in **(C)** MSK pan-cancer cohort and **(D)** ten human cancer types.

**Supplementary Figure 5. Disruption of *Mll3* reduces p16$^{Ink4a}$ and p19$^{Arf}$ protein levels and histone marks at their promoters in murine liver cancer.**

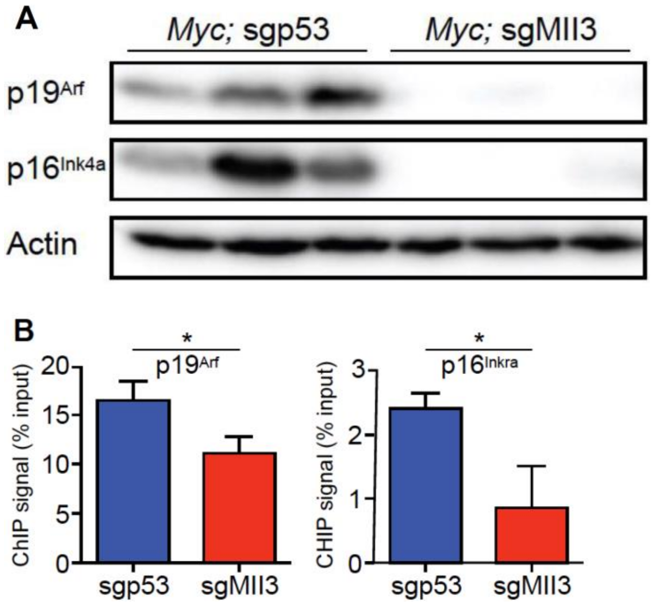


**(A)** Immunoblots for p16$^{Ink4a}$ and p19$^{Arf}$ proteins in three independent *Myc; sg*Mll3 and *Myc; sg*p53 HCC lines. β-actin was used as loading control. **(B)** ChIP-qPCR validation of H3K4me3 signals at *Arf* and *Ink4a* promoters. Values are mean ± SD from three independent *Myc*; sgMll3*;* and *Myc*; sgp53 liver tumor-derived cell lines. * = $P < 0.05$ (unpaired two-tailed t-test).

**Supplementary Figure 6. Mll3 directly regulates *Cdkn2a* expression in liver cancer cells.**

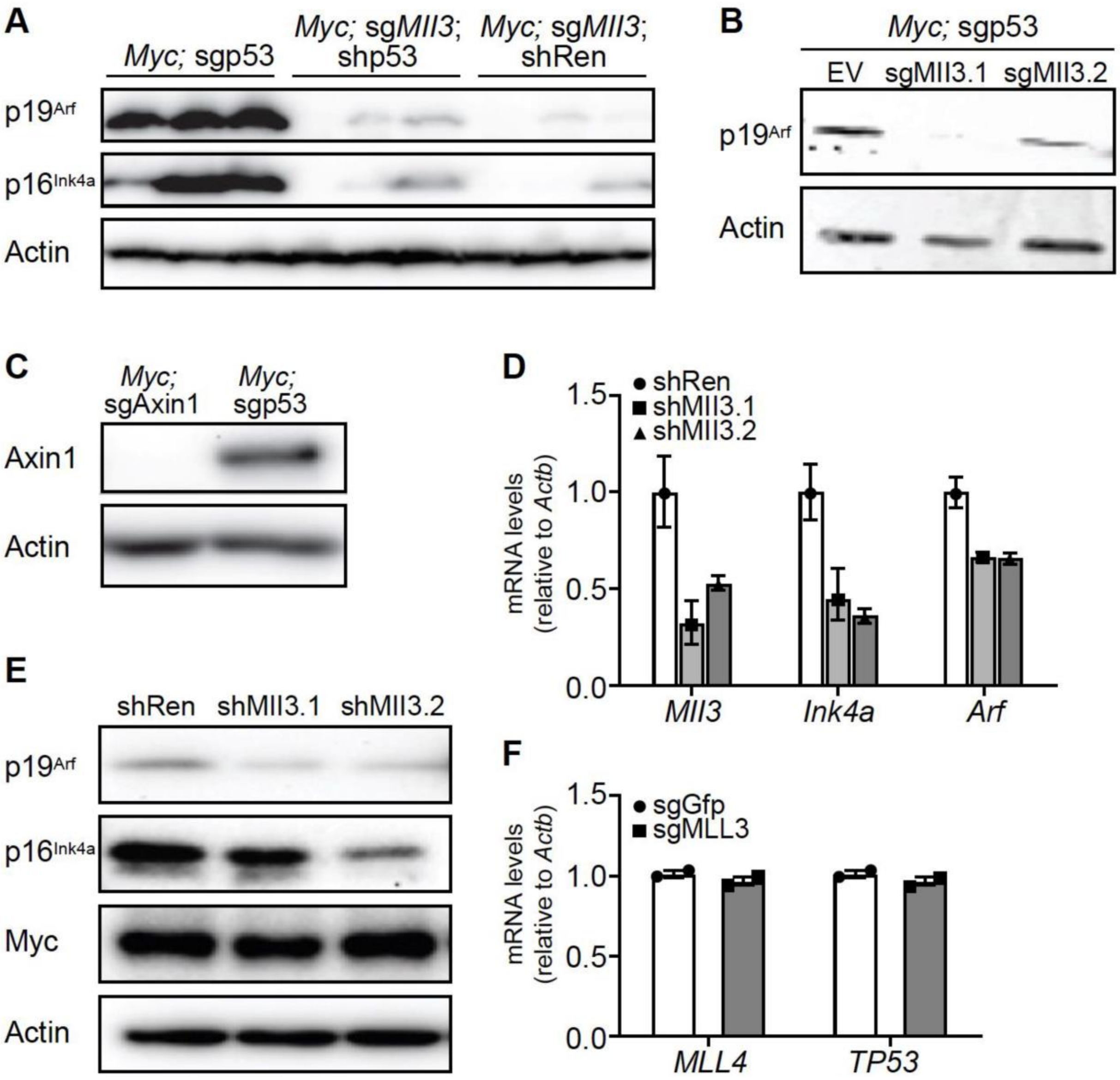


**(A)** Western blot analysis of p16$^{Ink4a}$ and p19$^{Arf}$ proteins in three independent *Myc*; sgp53 and *Myc*; sgMll3 cell lines expressing either a p53 shRNA (shp53) or a control shRNA (shRen). **(B)** Western blot analysis of p19$^{Arf}$ in *Myc*; sgp53 cell lines infected with two independent *Mll3* sgRNAs (sgMll3.1, sgMll3.2) or an empty vector control (EV). (**C**) Western blot analysis of Axin1 in *Myc*; sgAxin1 and *Myc*; sgp53 cell lines. **(D)** qPCR analysis of *Mll3*, *Arf* and *Ink4a* in Myc; sgAxin1 cell lines expressing two independent *Mll3* shRNAs or a control shRNA. Values are mean ± SD from experimental triplicates. **(E)** Western blot analysis of p16$^{Ink4a}$, p19$^{Arf}$ and Myc in *Myc*; sgAxin1 cell lines stably expressing two independent shRNAs targeting *Mll3* or shRen as a control. For all Western blots, β-actin was used as loading control. **(F)** qPCR analysis for mRNA expression of *MLL4 and TP53* in human HLE cells with sgGFP (control) or sgMLL3. Each data point represents the average of technical replicates (n = 3). Data are shown as mean ± SEM.

**Supplementary Figure 7. *Mll3* suppression reduces cell clearance upon enforced myc expression *in vivo.***

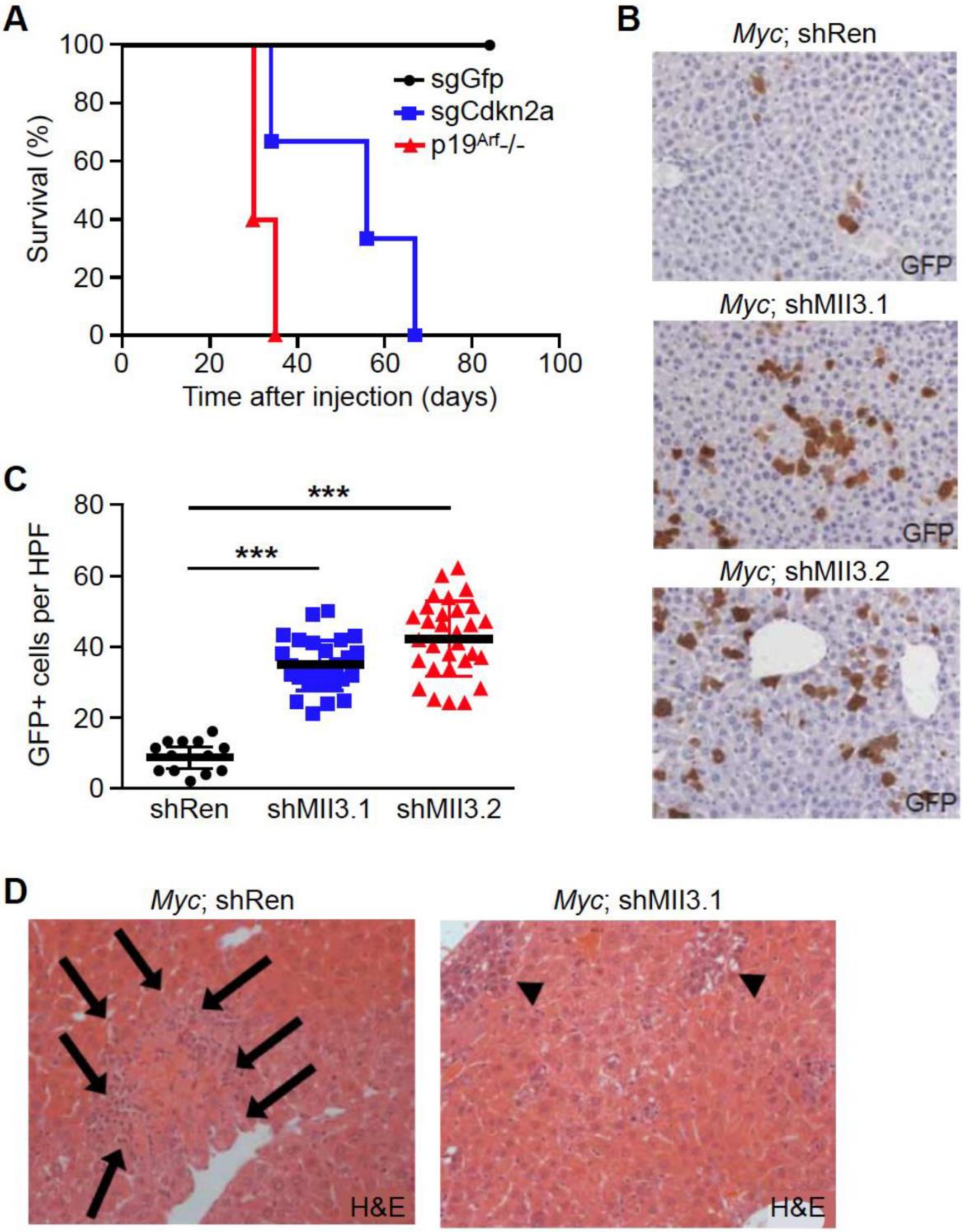


**(A)** Survival curve of wild-type mice injected with *Myc* transposon and pX330 expressing sgRNAs targeting *Cdkn2a* (*Myc;* sgCdkn2a, n = 5), or *p19^Arf^* -/- mice injected with *Myc* vectors (*Myc*; *p19^Arf^* -/-, n = 5). Wild-type mice injected with *Myc;* sgGFP (n = 3) served as a control. **(B)** Representative images of GFP-expressing cells detected by immunohistochemistry in murine livers 10 days after hydrodynamic injection of vectors enforcing stable expression of *Myc* and GFP-linked, constitutive shRNAs targeting *Mll3* or Renilla luciferase. **(C)** Quantification of GFP-positive cells in murine livers 10 days after hydrodynamic injection of indicated vector combinations. Data points represent GFP-positive cells in 10 different high-power fields (HPF) in three independent murine livers. **(D)** H&E images of murine livers receiving *Myc*; shRen or *Myc*; shMll3 vectors 5 days after injection. Arrows indicate apoptotic cell clusters. Arrow-heads indicate incipient neoplastic cell clusters.

**Supplementary Figure 8. Endogenous *Mll3* restoration triggers apoptosis and is accompanied by increased *Cdkn2a* expression.**

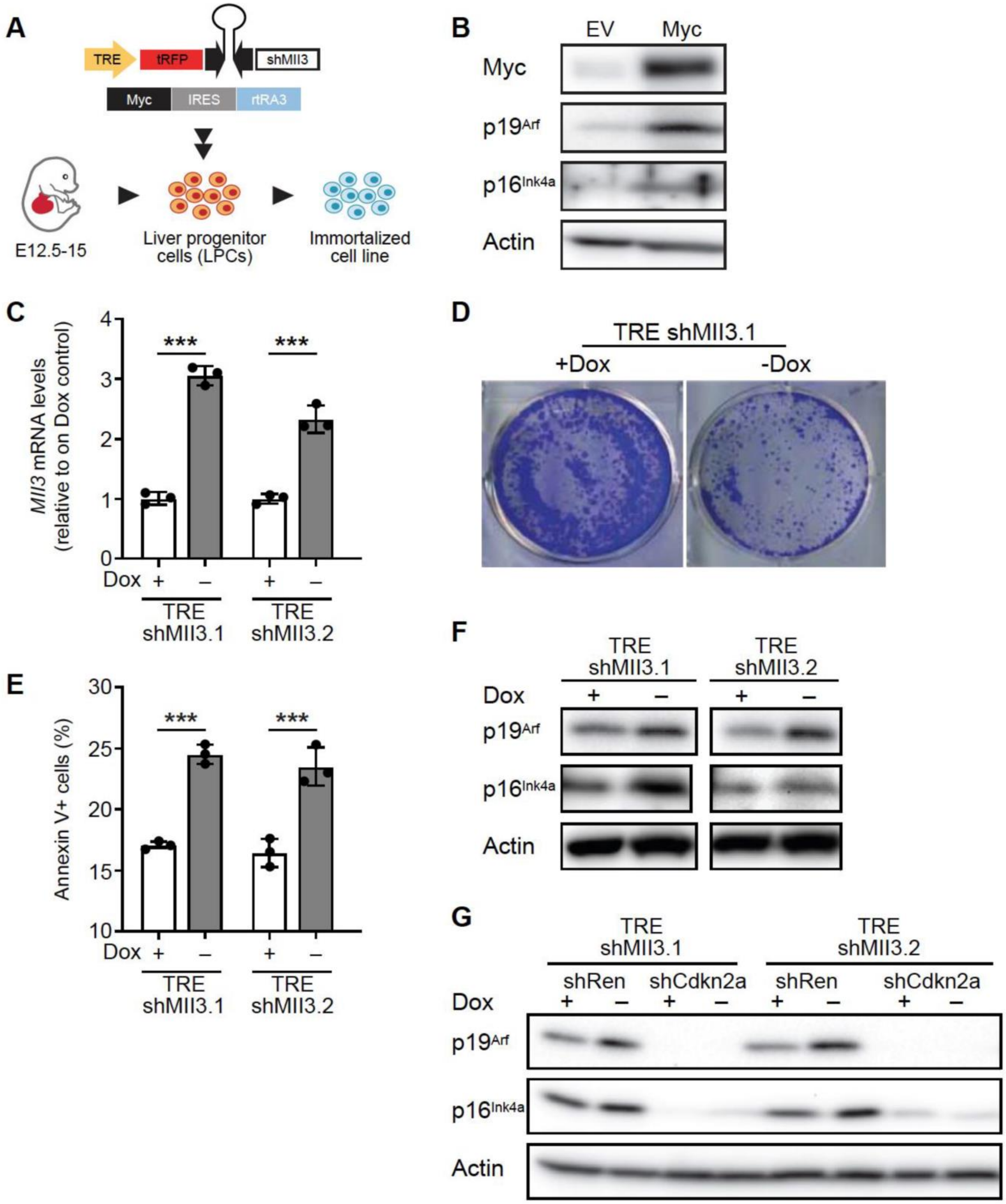


**(A)** Schematic for generating immortalized liver progenitor cell (LPC) lines from ED13 liver progenitor cells. **(B)** Western blot analysis of p16$^{Ink4a}$, p19$^{Arf}$ and MYC in wildtype LPCs 2 days after *Myc* transduction.

(**C**) *Mll3* mRNA expression before (on) and after (off) *Mll3* restoration in LPCs expressing with two independent tet-regulated *Mll3* shRNAs. Values are mean ± SD from technical replicates (n = 3). *** = $P < 0.001$ (unpaired two-tailed t-test). **(D)** Representative images of colony formation assay of LPCs four days after *Mll3* restoration. **(E)** Annexin-V labeling in *Myc*-rtTA3; TRE-shMll3 cells grown for 3 days with or without dox. Data are shown as mean ± SD of wells analyzed in triplicate. *** = $P < 0.001$, ** = $P < 0.01$ (unpaired two-tailed t-test). **(F)** Western blot analysis of $p16^{Ink4a}$ and $p19^{Arf}$ days after dox withdrawal in *Myc*-rtTA3; TRE-shMll3 cells. **(G)** Western blot analysis of $p16^{Ink4a}$ and $p19^{Arf}$ proteins in *Mll3* restored LPCs (two independent shRNAs targeting Mll3) with constitutive expressed control shRNAs (shRen) or *Ink4a*/*Arf*-targeting shRNAs (shInk4a/Arf). For all Western blots, β-actin was used as loading control.

### Supplementary Table S1

sgRNA sequences

| | |
|---|---|
| sgp53 (mouse) | ACCCTGTCACCGAGACCCC |
| sgMll3.1 (mouse) | AATCAGTGCCAACCAATGGC |
| sgMll3.2 (mouse) | CAGGTTGAAGACAAAAAGTA |
| sgChr8 (mouse) | ACATTTCTTTCCCCACTGG |
| sgGfp | GAATAGCTCAGAGGCCGAGG |
| sgMLL3 (Human, for CRISPRA) | CCCGCGCGTCAGGCCCGTC |

shRNA sequences

| | |
|---|---|
| shRenilla | AGGAATTATAATGCTTATCT |
| shMll3.1 (mouse) | GGAGACAAATATGTAGAGTT |
| shMll3.2 (mouse) | ACCAGTGATCACTTTACTAA |
| shCdkn2a (mouse) | AACACAAAGAGCACCCAGCGG |

qPCR sequences

| | |
|---|---|
| Actb (mouse)-Forward | GGCTGTATTCCCCTCCATCG |
| Actb (mouse)-Reverse | CCAGTTGGTAACAATGCCATGT |
| p16-Ink4a (mouse)-Forward | CCATCTGGAGCAGCATGGAGT |
| p16-Ink4a (mouse)-Reverse | ACGTGAACGTTGCCCATCATC |
| p19-Arf (mouse)-Forward | CGCTTCTCACCTCGCTTGTC |
| p19-Arf (mouse)-Reverse | CAGTGACCAAGAACCTGCGA |
| Mll3 (mouse)-Forward | CAGGAGGGCCTGCAAGATAC |
| Mll3 (mouse)-Reverse | TATCCTCCGGTTGGAGCTGA |
| ACTB (Human)-Forward | AAGAGCTACGAGCTGCCTGA |
| ACTB (Human)-Reverse | TCCATGCCCAGGAAGGAAGG |
| p16-INK4A (Human)-Forward | GAGCAGCATGGAGCCTTCGG |
| p16-INK4A (Human)-Reverse | TGGATCGGCCTCCGACCGTAA |
| p14-ARF (Human)-Forward | GCAGGTTCTTGGTGACCCTC |
| p14-ARF (Human)-Reverse | TAGACGCTGGCTCCTCAGT |
| MLL3 (Human)-Forward | GAAACGCTGTAGCCTGTCCT |
| MLL3 (Human)-Reverse | CCCTGAGTCTCCTTTGGCAG |
| MLL4 (Human)-Forward | GCCCTTTCTTCAAGGTGGACT |
| MLL4 (Human)-Reverse | CGGGTTCCGGGCTAAAGAAG |
| TP53 (Human)-Forward | TGACACGCTTCCCTGGATTG |
| TP53 (Human)-Reverse | TCATCCATTGCTTGGGACGG |

ChIP-qPCR sequences

| | |
|---|---|
| p16-Ink4a promoter (mouse)-Forward | GATGGAGCCCGGACTACAGAAG |
| p16-Ink4a promoter (mouse)-Reverse | CTGTTTCAACGCCCAGCTCTC |
| p19-Arf promoter (mouse)-Forward | GACCGTGAAGCCGACCCCTTCAGC |
| p19-Arf promoter (mouse)-Reverse | GGGGTCGCTTTCCCCTTCGG |
